# Active subspace learning for coarse-grained molecular dynamics

**DOI:** 10.1101/2025.10.13.682174

**Authors:** Anna Wojnar, Stephen Pankavich, Alexander J. Pak

## Abstract

We introduce Active Subspace Coarse-Graining (ASCG), an interpretable framework for systematic bottom-up coarse-graining trained from atomistic molecular dynamics simulations that simultaneously defines the coarse-grained mapping, effe
ctive interactions, and the equations of motion within one unified mathematical framework. We employ active subspace learning to identify linear projections of atomistic degrees of freedom that maximally describe gradients of the potential energy, yielding a reduced set of coarse-grained variables that capture the dominant collective motions across the potential of mean force. Effective coarse-grained forces and noise terms are obtained directly from the same projection, eliminating the need for separate parameterization schemes. We demonstrate the ASCG method on three biomolecules: dialanine, Trp-cage, and chignolin. We show that free energy surfaces are recapitulated with Jensen-Shannon divergences as low as 0.034 while eliminating all solvent degrees of freedom and reducing solute dimensionality by more than 90%. The ASCG trajectories are integrated with timesteps up to 100 fs, around four to ten times larger than those possible with conventional coarse-graining methods, while ASCG models remain accurate with as little as 100 ns of training data. These results establish ASCG as a robust, data-efficient approach for learning complete coarse-grained representations directly from molecular forces, while representing a departure from traditional particle-based models.

## Introduction

Classical molecular dynamics (MD) simulations are a key tool for molecular modeling in numerous disciplines, such as for the study of protein folding^1-3^ and ligand binding^4-7^ in biological sciences. All-atom (AA) MD simulations entail representing every atom in a system as a particle and integrating their motions over time via Newtonian mechanics or related equations of motion. This approach offers researchers insight into the dynamics of (biomacro)molecular systems on atomic scales. While there have been advancements in experimental techniques, such as in single-particle fluorescence microscopy, to probe dynamical behavior on atomic scales, these methods often require perturbations (such as functionalized probes) that may alter the system dynamics or are limited in the temporal resolution (on the order of ms) that is able to be imaged.^8, 9^ The advantage of AAMD simulations in probing molecular systems at such fine spatiotemporal resolutions is why the literature has seen an increase in AAMD simulations from both computational and experimental molecular biologists.^10^ However, the scale of systems accessible to AAMD are inherently bound by computational limitations.

Even with the growing throughput of supercomputers, achievable timescales for AAMD simulations are on the order of microseconds (or milliseconds using specialized hardware^11-13^), with this achievable timescale decreasing as the system size increases.^14^ However, many types of behaviors of interest, such as protein conformational switching or assembly, require inaccessible timescales on the order of milliseconds and beyond (even as large as hours to days).^15^ Much of this limitation arises from the integration timestep, which is limited to 1-2 fs.^10, 16^ The choice of timestep is limited by the fastest modeled motions, such as the high-frequency vibrational modes associated with bond stretching.^17^ Using a larger timestep introduces numerical integration errors, which accumulate over the course of the simulation and often render a simulation either unstable or inaccurate.^17^ To probe behavior at longer timescales, numerous techniques have been introduced, including enhanced sampling and coarse-graining, the latter of which is the focus of this paper.

Coarse-grained (CG) modeling involves the conversion of AA models into lower resolution forms by: (1) mapping groups of atoms into individual CG sites, sometimes excluding other atoms (such as solvent or ions), and (2) defining the effective interactions between CG sites.^18-20^ The resultant CG model reduces the degrees of freedom of the system by orders of magnitude, enabling simulations of larger spatial scales compared to AAMD at comparable computational cost. Another advantage is the ability to increase the simulation timestep as the mapping eliminates high-frequency motions and effectively smooths the underlying energy surface. However, the benefits of CG models come with a tradeoff, as accuracy is sacrificed in comparison to the higher-resolution AA models. The accuracy of the CG model is contingent upon the strategies used to create the model, which are broadly defined by two categories: top-down and bottom-up CG models. Top-down models typically map CG sites based on chemical intuition, then tune CG interaction potentials to reproduce macroscopic properties (usually obtained from experiments)^21-25^, but, as a result, tend to be inaccurate with respect to microscopic correlations^26-28^. Conversely, bottom-up approaches aim to systematically reproduce microscopic correlations (obtained from higher resolution data, such as from AAMD) but provide no guarantee that macroscopic properties are preserved.^19, 20, 28-32^ Nonetheless, cases in which bottom-up models lack macroscopic consistency are meaningful; the implication being that the enforced microscopic correlations are either incorrect or incomplete. One can alternatively view bottom-up methodologies as a hypothesis testing framework to identify the most important microscopic aspects that are central to emergent macroscopic behavior.

Within the field of bottom-up CG modeling, numerous methodologies have been proposed for both mapping and parameterization. To date, mapping approaches include center-of-mass^33^, center-of-charge^34^, or clustering algorithms^35, 36^ amongst others. Algorithms to identify and compute effective CG interactions include force-matching^33, 37-40^, relative entropy minimization^41^, and energy-matching^42^ amongst others, while a growing number of machine learning based approaches^31, 43-48^ have been introduced to augment the basis sets used for these methods. While the mapping and parameterization steps are inherently coupled, i.e., the fidelity of the interaction potential is restricted by the mapping, most mapping and interaction parameterization methodologies are investigated independently; in fact, the choice of appropriate mapping and parameterization method remains an open question and is likely problem- and system-specific. To our knowledge, there have only been a few attempts to combine mapping and parameterization in a unified framework. Wang and Bombarelli^46^ introduced a variational auto-encoder (VAE), a generative machine learning model, to learn a latent (lower-dimensional or CG) representation of AA data, which was additionally conditioned on atomic forces inspired by the force-matching strategy of Voth and coworkers^33, 37-40^. Yet, perhaps as a result of the discrete CG mapping enforced by the VAE, even the simple gas molecules and alkanes that were tested required between a 1:1 to 3:1 ratio mapping of heavy atoms to CG sites to maintain accuracy, which limits the potential gains by the CG model. More recently, the use of VAEs as a nonlinear “mapping” function was also investigated by Ferguson and coworkers^49, 50^ and Koumoutsakos and coworkers^51^, although rather than learning the effective CG interactions, these authors learned approximate functions to propagate the dynamics of the latent variables directly. However, the interpretability of the mapping (and the effective interactions) from these latter cases is made difficult by the complexity of the trained neural networks.

In this manuscript, we introduce a linear and interpretable methodology for bottom-up CG modeling of both the mapping and parameterization function. Our central idea is to identify linear projections of AA degrees of freedom as both CG configurational variables and their associated forces. The projections are identified through active subspace learning, which is a dimension reduction technique that discovers the directions of an input space that maximally describe gradients of an output scalar quantity of interest.^52-56^ We show that by learning the active subspace based on the potential energy as the output quantity of interest, we implicitly learn a set of CG variables and the associated potentials of mean force (PMFs) along these variables. Our methodology, which we call Active Subspace Coarse-Graining (ASCG), is demonstrated for three biomolecules: dialanine, chignolin, and Trp-cage.^57, 58^ Our analysis reveals two notable advantages of ASCG compared to traditional CG modeling strategies: (1) we remove *ad hoc* decisions associated with separate methods for mapping and parameterization and (2) we enable integration timesteps that exceed even those made possible by CG models. Finally, we show that training ASCG models is inherently data efficient, requiring as little as 100 ns of AAMD data for the three tested biomolecules.

### Theory and Methods

#### Theory

Active subspace methods^52, 54, 59^ represent a collection of recently developed dimension reduction techniques that identify important directions, the span of which is referred to as the *active subspace*, in a high-dimensional space of input variables ***x*** and their influence on an associated output quantity of interest, *g*(***x***). The identification of these subspaces allows for (i) global sensitivity measurements of outputs with respect to model inputs and (ii) the construction of reduced-order, surrogate models that greatly decrease the dimensionality of the input parameter space.^55, 60-64^

In the context of MD simulations, we propose that active subspace methods will allow us to reduce the complexity of expensive AA MD simulations via an active-subspace-informed surrogate model, which will serve as a CG model, i.e., the aforementioned ASCG model. Here, we define the vector of input variables ***x*** ∈ ℝ ^3*n*^ as the atomistic configurational degrees of freedom for *n* atoms. For simplicity, we use the 3*n* Cartesian coordinates of the biomolecule of interest after remapping to a local coordinate system (see *Supplementary Information*) in order to maintain roto-translational invariance, noting that six of those degrees of freedom will always be zero; we refer to these as local frame transformation (LFT) features. Note that we discard all solvent degrees of freedom, yet the influence of solvent remains implicit in the evolution of the solute degrees of freedom. We also specify that the output quantity of interest is the potential energy *E*(***x***), i.e., *g*(***x***) = *E*(***x***), which can also be a function of any related representation of ***x***. Consequently, the active subspace can be thought of as a low-dimensional projection of configuration space that maximally represents the largest variability (or gradients) of potential energy.

To clarify the meaning behind the active subspace, we will describe the standard formulation of the gradient-based active subspace method^52^ adapted to data readily available from AAMD simulations. Let ***x***_***m***_ ∈ ℝ ^3*n*^ denote a vector of mass-weighted positions ***x*** for each independent variable *x*_*i*_ and *x*_*m,i*_ (with *i* = 1,…,3*n* ):

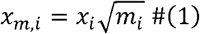

where *m*_*i*_ is the mass associated with the atom indexed at *i*. We perform mass-weighting to simplify the transformation of the equations of motion, as discussed later. Then, let 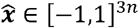 denote a vector of standardized ***x***_***m***_ values, where we assume that the mass-weighted position variables have been shifted and scaled so that they linearly scale from -1 to 1. Here, each variable 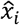 is independently obtained via:

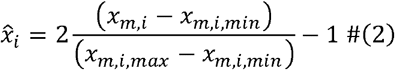

where *x*_*m,i,min*_ and *x*_*m,i,max*_ represent the minimum and maximum values of *x*_*m,i*_, respectively, from the AAMD trajectories. Using the chain rule, we define the vector of gradients to the output quantity with respect to 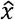 as 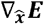 via:

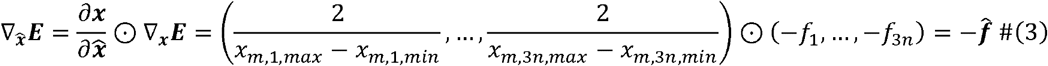

where *f*_*i*_ refers to the force acting on degree of freedom *i*, ⊙ is the Hadamard product, and 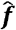 is the effective force along 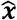. We emphasize that the forces ***f*** ∈ ℝ ^3*n*^ are computed directly during AAMD simulations and thus require no extra computation to obtain.

Next, the covariance matrix is defined as:

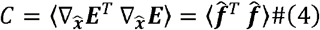

where ⟨ ⟩denotes an ensemble average. Since AAMD simulations already sample from the underlying thermodynamic ensemble, we compute *C* via:

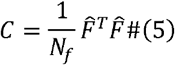

where *N*_*f*_ is the total number of simulation frames and 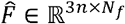 is the matrix of effective forces 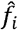 across all simulation frames. Note that *C* ∈ ℝ ^3*n* × 3*n*^ is a symmetric, positive semi-definite matrix, with the spectral decomposition^65^:

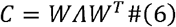

where *W* is an orthogonal matrix whose columns *w*_*i*_ are orthonormal eigenvectors of *C* and Λ = diag (*λ* _1_,… *λ*_3*n*_) is the diagonal matrix of eigenvalues of *C*. Here, the eigenvalues in Λ are listed in descending order, i.e., *λ* _1_ ≥*λ* _2_ ≥ … ≥ *λ*_3*n*_ ≥0, and the associated eigenvectors within the same column are arranged with their corresponding eigenvalues. Crucially, *λ*_*i*_ measures the average gradient in 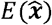 (or force) subject to perturbations in 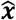 along *w*_*i*_, as they are related by the identity:

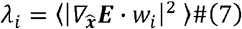

for *i* =1,…,3*n* . For example, if λ_i_ = 0, then Eq. (7) shows that 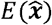 is constant along the direction *w*_*i*_, and such directions can be ignored when studying the behavior of 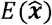 under changes to the input space 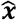. Conversely, if *λ*_*i*_ is large, then we deduce from Eq. (7) that 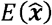 changes considerably in the direction of *w*_*i*_.

Now, suppose that a spectral gap exists that separates the first *m*< 3*n* eigenvalues from the trailing 3*n* − *m* eigenvalues, as the former are significantly larger. Then, the eigenpairs can be block separated as:

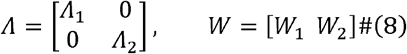

where Λ_1_ ∈ ℝ^*m* × *m*^ contains the “large” eigenvalues of *C*, Λ_2_ ∈ ℝ ^(3*n* − *m*) × (3*n* − *m*)^ contains the “small” eigenvalues, and *W*_*k*_ contains the eigenvectors associated with each Λ_*k*_ for *k* =1,2. The spectral gap can be determined by logarithmic plotting or using relative differences to estimate the amount of variability retained by each eigenvalue.^55^ This gap will correspond to differences of around an order of magnitude and thus allow one to compartmentalize the greatest eigenvalues within Λ_1_ and the remaining (lesser) eigenvalues in Λ_2_.

With the decomposition in Eq. (8), we can represent any sample 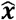 of the input space by:

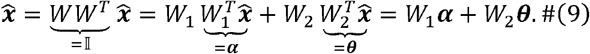

Evaluating the output *E* at 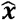 is equivalent to doing so at *W*_1_***α*** + *W*_2_ ***θ***, and we can further approximate 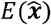 using:

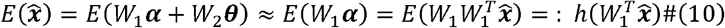

where *h* is the projection of *E* onto the range of *W*_1_, and 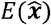 simplifies to 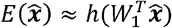. Due to the small eigenvalues corresponding to *W*_2_, Eq. (7) shows that changes to ***θ*** will have only extremely small influence on the values of 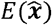. Therefore, *W*_1_ ∈ ℝ^3*n*×*m*^ is defined to be the *active subspace* of the model, while *W*_2_ ∈ ℝ^3*n* × (3*n* − *m*)^ is the *inactive subspace*. The linear combinations that generate these subspaces represent the contributions of differing 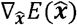 with respect to input variations. We emphasize that while the described dimension reduction bears some similarity to principal component analysis (PCA)^66^, the active subspace method is a supervised dimension reduction technique whereas PCA is an unsupervised technique that has no knowledge of 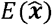.

With the active subspace defined, we now discuss the formulation of the CG (or reduced) model, hereafter referred to as the ASCG model. In a typical CG model, one first defines the mapping operator from AA to CG resolution.^33-36^ Then, the effective CG interactions are determined using several available methods.^33, 42, 43, 67, 68^ Finally, the CG model is integrated using standard equations of motion. Here, we show that the active subspace formulation can be leveraged to simultaneously define the CG mapping, effective interactions, and equations of motion in a unified manner.

The CG mapping is defined by the linear projection shown in Eq. (9). Specifically, we refer to the set of variables ℝ ^*m*^ as the *m* ASCG degrees of freedom:

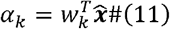

with *w*_*k*_ representing the th (with *k* =1,…,*m* ) column of _1_. With this, it is typical to implement a nonlinear least-squares fit to approximate the function *h* defined in Eq. (10) to represent the effective ASCG interactions. However, 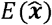 can be difficult to fit due to its noise. We therefore opted to determine the effective interactions by first addressing how the mapping in Eq. (11) affects the underlying fine-grained equations of motion.

Consider the Langevin equation of motion that describes the integration of variable *x*_*i*_ in the canonical (constant NVT) ensemble:

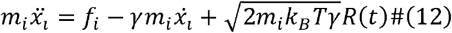

where 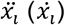 is the acceleration (velocity) of *x*_*i*_, *γ* is the friction (or damping) constant, *k*_*B*_ is the Boltzmann constant, *T* is the temperature, and *R*(*t*) is random white noise sampled from a Gaussian distribution with zero mean and unit variance. Applying the mass-weighting and standardization described in Eq. (1) and (2) to Eq. (12), we rewrite the equation of motion as:

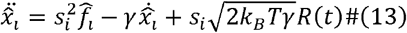

which describes the Langevin equation of motion for the standardized variable 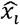 where 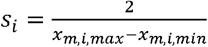. Next, we consider the set of Langevin equations of motion for 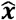 and apply the projection by *W*_1_ such that:

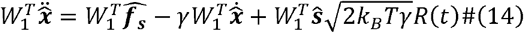

where is the vector of scaled forces with element and is a vector of elements . Applying the definition from Eq. (11) and by analogy to Eq. (12), we rewrite Eq. (14) to find the set of ASCG equations of motion:

where ( ) is the vector of accelerations (velocities) of, is the vector of effective forces acting on, and is the vector of standard deviations for the random noise. Therefore, in lieu of approximating the function as an estimator for, we directly compute the effective force for each ASCG variable as:

while

completes the definition of the ASCG equations of motion.

The ASCG mapping (Eq. (11)), effective forces (Eq. (16)), and equations of motion (Eq. (15)) demonstrate the unified approach to coarse-graining via active subspace projection, as also depicted in **Figure 1**. Below, we demonstrate our approach on three biomolecular systems: dialanine, chignolin, and Trp-cage. We also note that other geometric degrees of freedom can be used to describe, such as pairwise distances commonly seen in AA and CG force-fields^33, 67, 68^. The framework described above can be remapped to these variables through application of the chain rule, which we leave for a future publication.

**Figure 1.**
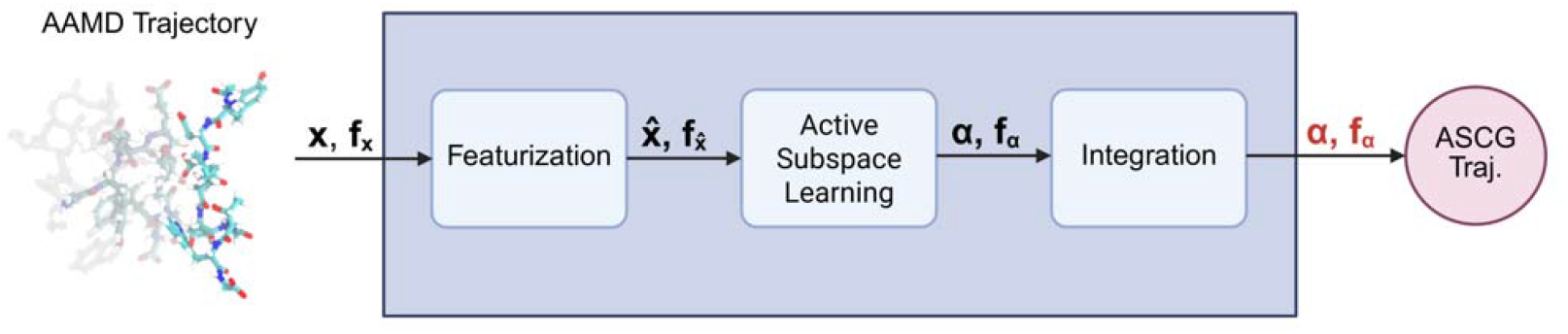
Active subspace coarse-graining (ASCG) framework. Schematic of the proposed ASCG methodology. All-atom molecular dynamics (AAMD) simulations, consisting of positions ( ) and their corresponding forces (*f*_*X*_ ), are featurized to enforce roto-translational invariance and standardization. Active subspace learning is performed on the featurized positions 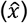 and forces 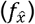, where we identify the directions in 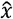 space that most influence changes to the potential energy. Applying dimensional reduction with active subspace learning, we identify a mapping and parameterization of 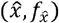 into ASCG variables (*α, f*_*α*_ ). Using ASCG equations of motion, we integrate dynamics to generate an ASCG trajectory, which we then analyze to compare with AAMD statistics.

#### All-Atom Molecular Dynamics

Atomic structures for Trp-cage (PDB 1L2Y^58^) and chignolin (PDB 1UAO^57^) were obtained from the Protein Data Bank. The atomic structure for dialanine was constructed using the Molefacture plugin in Visual Molecular Dynamics (VMD) 1.9.4.^69^ All MD simulations were performed using GROMACS 2023.2^70^, using the CHARMM36m forcefield and modified TIP3P water.^71^ Each biomolecule was placed in a cubic box with a buffer distance of 2 nm on all sides. All biomolecules were simulated in aqueous solution of 150 mM NaCl. The dialanine, Trp-cage, and chignolin systems contain a total of 27,062, 14,467, and 14,449 atoms, respectively (23, 304, and 138 atoms for solute only). Energy minimization was performed using steepest descent until the maximum force was lower than 500 kJ/mol/nm. Equilibration was performed in three stages: (i) first, water and ions were relaxed in the constant NVT ensemble at 300, 270, or 420 K (for dialanine, Trp-cage, and chignolin, respectively) for 10 ns using the stochastic V-rescale thermostat^72^ with 0.1 ps damping time, (ii) next, all atoms were relaxed in the constant NPT ensemble at the same temperatures and 1 bar for 30 ns using the same thermostat and C-rescale barostat^73^ with 0.5 ps and 5.0 ps damping time, respectively, and (iii) finally, all atoms were relaxed in the constant NVT ensemble at the same temperatures for 10 ns using the same thermostat with 2.0 ps damping time; all simulations used a 2 fs timestep with LINCS restraints^74^ for bonds containing hydrogen and Particle Mesh Ewald^75^ for long-range electrostatics.

For dialanine, data was subsequently collected over 1 µs in the constant NVT ensemble at 300 K across four independent replicas. The positions and forces of the peptide were saved every 10 ps for each replica (i.e., 400k samples total). For the more complex Trp-cage and chignolin cases, data was collected using two steps. First, a long trajectory was run in the constant NVT ensemble for 500 ns. Throughout the long trajectory, the simulation state was saved every 5 ns and each state was used to run short simulations (a total of 100 simulations). In the case of Trp-cage, short simulations were performed in the constant NVT ensemble at 270 K with protein configurations and forces saved every 2 ps over 20 ns per short trajectory. For chignolin, the same general procedure was used but simulated at 420 K and data saved every 1 ps over 10 ns per short trajectory.

#### Coarse-Grained Mapping

As a pre-processing step before featurization of the positions and forces, an intermediate CG mapping was performed to reduce dimensionality prior to active subspace learning. All mapping procedures used center of mass mapping as implemented by OpenMSCG 0.9.0.^76^ For dialanine, we used a united atom representation, where a CG site consists of each heavy atom and any bonded hydrogen atoms. For Trp-cage and chignolin, we mapped the backbone and sidechain of each amino acid as two separate CG sites; glycine was mapped as one CG site.

#### Active Subspace Coarse-Graining

The workflow as described above (and in **Figure 1**) was implemented in Python using NumPy.^77^ Before calculation of the covariance matrix (Eq. 5), the six input features that are always zero (due to translation and rotation into the local coordinate system) were filtered out as their presence led to high condition numbers and the eigenvector components corresponding to these six features had negligible magnitudes.

To perform multivariate interpolation of computed ASCG forces, we used k-nearest neighbor regression as implemented in *scikit-learn*.^78^ For each biomolecule, the projected ASCG forces from AAMD simulations were averaged into *m*-dimensional bins of width 0.02 for dialanine and trp-cage, and 0.03 for chignolin, where *m* is the number of ASCG components for that system. The average force of each bin was used to train the interpolation scheme. Interpolation was implemented with a weighted distance between the 50 nearest neighbors for dialanine, and 500 nearest neighbors for trp-cage and chignolin, using the KDTree algorithm^79^ with a leaf size of 25 to detect neighbors. For values outside of the range of sampled ASCG positions, a fill value was set to the same order of magnitude as the ASCG forces at each endpoint.

We implemented ASCG simulations via integration of Eq. 15 using NumPy. Each biomolecule was simulated at the same temperature as their respective AAMD simulation. We performed 100 independent replicas per system at 50 ns per replica; initial conditions were generated by selecting 100 random frames from the AAMD trajectories for coordinates; initial velocities were calculated using central finite differencing. Dialanine was simulated using a timestep of 50 fs and a friction constant of 0.5 ps^-1^. Both Trp-cage and chignolin were simulated with a timestep of 100 fs and a friction constant of 0.5 ps^-1^. The Jensen-Shannon divergence (JSD) was computed for the *m*-dimensional probability density over ASCG variables between that of the AAMD reference and simulated ASCG trajectories.

To test data scarcity effects, we incrementally truncated the trajectories used for ASCG model training. For dialanine, each of the four independent replicas were truncated by the same amount. For Trp-cage and chignolin, as each replica was seeded from different points along the initial long trajectory, trajectories past specified time points were discarded. ASCG models were retrained using the truncated datasets and the same hyperparameters for interpolation. The same integration hyperparameters were used for ASCG simulations. We noticed that for some truncated datasets, the AAMD probability density was rotated in reference to the full-length AAMD probability density (**Fig. S1**), artificially inflating the calculated JSD between the ASCG trajectory and full-length AAMD reference. To address this, before computing JSDs between the ASCG trajectory and the full-length AAMD reference, we corrected this rotation (details in *Supplementary Information*).

## Results and Discussion

### Proof of Concept: Dialanine

We first tested the ASCG workflow on dialanine given its relative simplicity. Briefly, we performed AAMD simulations of dialanine and then instituted united atom mapping of dialanine (**Figure 2A**), a high-resolution CG mapping wherein heavy atoms are grouped with their bonded hydrogens. Next, we prepared LFT features for both positions and forces and then rescaled them as described in *Theory and Methods*.

**Figure 2.**
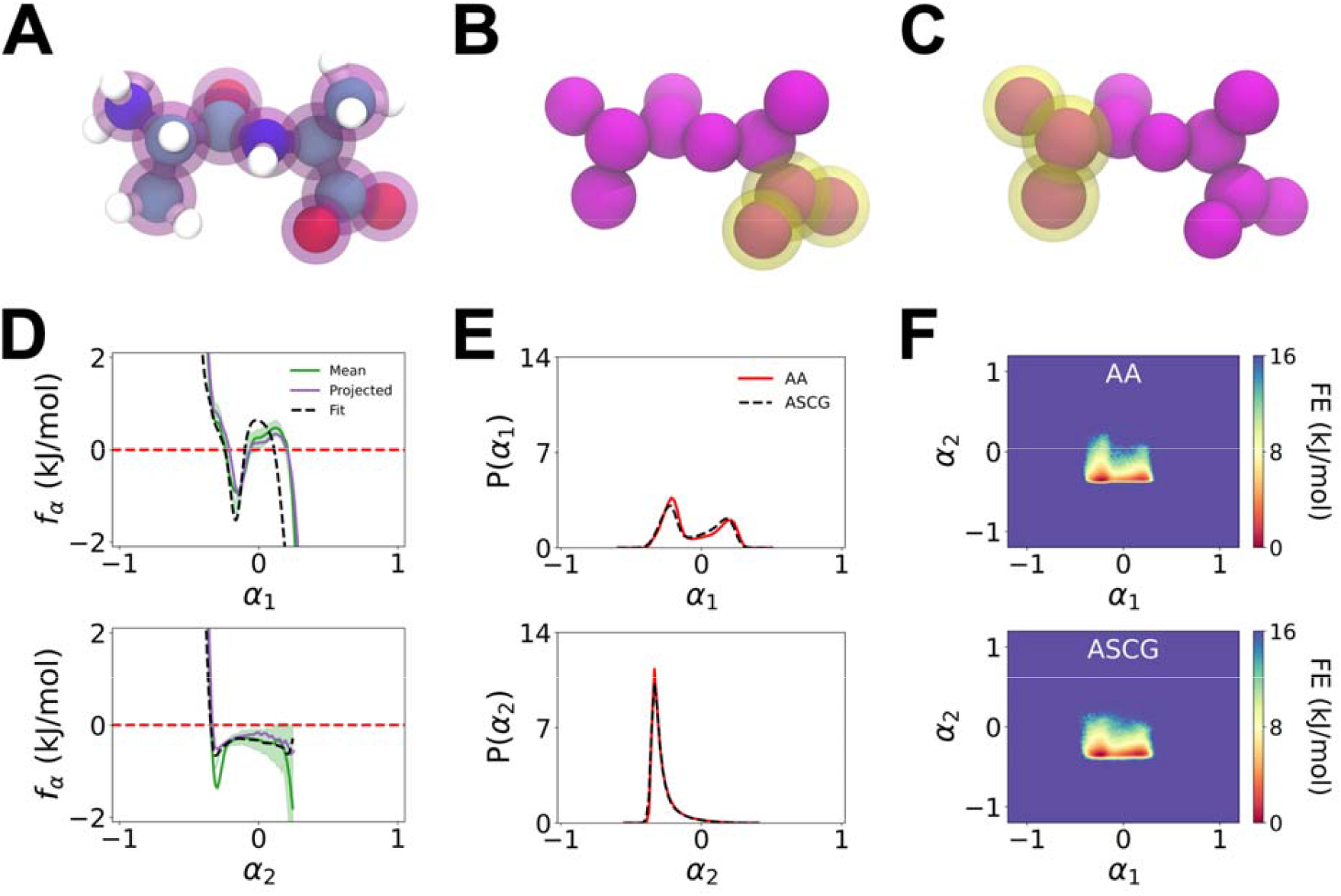
ASCG mapping, parameterization, and integration results for dialanine of the first two components. (A) Schematic of united atom mapping for dialanine, where the 11 united atom sites are depicted in purple. (B-C) The ASCG mapping of (B) and (C) is depicted in yellow, where and represent the C-terminus and N-terminus, respectively. (D) Force ( ) profiles for (top) and (bottom). The mean force represents the average forces along each ASCG component after transformation, shown with a 95% confidence interval. Projected forces represent forces estimated by calculating a 1D free energy surface from the 1D probability density of the atomistic data and then taking the negative gradient. Fit forces are predicted forces after fitting a multivariate nearest neighbors algorithm to the averaged ASCG transformed forces. (E) 1D probability distributions along (top) and (bottom) comparing the atomistic (AA) distribution with the ASCG distribution calculated after integrating and . (F) 2D slice of the free energy (FE) surfaces of and generated from AAMD trajectories (top) and from ASCG integration of through (bottom). The JSD between the underlying 2D probability distributions of and between the AAMD surface and the ASCG-generated surface is 0.023.

The ASCG workflow identifies linear projections of the input positions that maximally describe the gradients to the output potential energy. We find that the top three ASCG components contain 43% explained variance in force (**Fig. S2A**). This is comparable to the explained variance of PCA trained on positions alone (43% for the top three components). The spectral ratio between adjacent components, which we define as drops significantly after three components (**Fig. S3A**) and justifies focusing on only the top three components. As a result, the three-component ASCG model represents a 91% reduction in dimensionality compared to the original LFT features.

The eigenvectors associated with the top three components encode both the ASCG mapping (via Eq. 11) and ASCG effective force (via Eq. 16). **Figures 2B-C** highlight the LFT positions with the largest contribution to the top two components (top three components in **Fig. S4**), respectively, where contribution is filtered by eigenvector component magnitudes larger than 0.1. We find the mapping physically meaningful as for *α*_1_ and *α*_3_, the respective first and third ASCG variables primarily capture the rotations of the carboxylate group in the C-terminus, and in the case of *α*_2_, capture the rotations of the methyl and amine groups in the N-terminus, respectively. We emphasize, however, that the mapping does not yield traditional particle-based CG sites. Instead, each of these ASCG variables can be interpreted as CG descriptors of collective motions, albeit ones that can be propagated with their own equations of motion. To propagate dynamics, we compute the mean effective ASCG forces as a function of *α*_1_ through *α*_3_, as seen in **Figure 2D** and **Fig. 4B**. Similar to conventional forces acting on inter-particle distances, we observe ASCG forces that become increasingly repulsive at low *α* values and increasingly attractive at high *α* values. The most important observation from **Figure 2D** is that the ASCG forces are also consistent with the negative gradient of the potentials of mean force (PMFs) for *α*, where the PMF is computed by −*k*_*B*_*T* ln (*P*(*α*)) and *P*(*α*) is the probability distribution of *α*. In other words, we have shown that the ASCG approach discovers a projection of atomistic forces that is consistent with the PMF describing the same projection of atomistic positions. This observation is one of the central benefits of the ASCG approach, and we reinforce this point with comparison to PCA (discussed later).

Next, we simulated the dynamics of *α*_1_ through *α*_3_ using the equations of motion described by Eq. 15. We integrated 100 independent ASCG trajectories of dialanine, each 50 ns with a timestep of 0.05 ps and *γ* = 0.5 ps^-1^; note that this *γ* value is equivalent to the damping time from the AAMD integrator, but the effective ASCG friction is subsequently modified as seen in Eq. 17. The resulting 1D probability densities and 2D slice of the free energy surfaces (FESs) depicted in **Figure 2E** show the qualitative similarity between the sampled AAMD and ASCG conformations, specifically the two major conformational states near (*α*_1_, *α*_2_ ) (-0.22, -0.35) and (*α*_1_, *α*_2_) (0.18, -0.35). The first state represents the configuration where the N-terminal amine and the C-terminal carboxylate are on the same side of dialanine, while in the second state the two moieties are on opposite sides. To quantify the similarity between the ASCG-generated and AAMD distributions, we calculated the 1D JSD of the three 1D distributions and the 3D JSD of the 3D distribution, where 0 indicates a perfect match between the ASCG and AAMD statistics, and larger JSD values indicate greater differences between the two. The excellent recapitulation of the AAMD FES by ASCG is reflected by the low 1D JSDs of 0.014, 0.006, and 0.014 for *α*_1_, *α*_2_ and *α*_3_, respectively, and 3D JSD of 0.034. Full results for the top three components can be found in **Fig. S4**. We found it necessary to use the three-body ASCG force approximated by our multivariate interpolator, i.e., a force that depends on *α*_1_, *α*_2_ and *α*_3_. If we treat the ASCG force as univariate and independent for each ASCG component, we find that the structure of the joint distribution (such as the existence and location of the metastable states) is lost (**Figs. S5, S6, and S7** for dialanine, Trp-cage, and chignolin, respectively). We found that for components that are not coupled, such as *α*_1_ and *α*_2_ or *α*_2_ and *α*_3_ for dialanine, sampling is not significantly impacted by treating the ASCG force as univariate and independent (**Fig. S5**).

In summary, we have demonstrated that ASCG is a viable workflow for all three phases (i.e., mapping, parameterization, and integration) of systematic coarse-graining. In the next section, we test the robustness of the ASCG workflow on more complex mini-proteins: Trp-cage and chignolin.

#### Extensions to Trp-cage and Chignolin

We now investigate the fidelity of the ASCG workflow on the fast-folding miniproteins Trp-cage and chignolin, two examples of increased complexity compared to dialanine. Trp-cage is a 20 residue miniprotein that maintains a stable folded structure consisting of an *α*-helix, 3_10_-helix, and polyproline II helix with a hydrophobic core.^58, 80^ Chignolin is a 10 residue miniprotein that forms a stable *β*-hairpin structure.^57^ Unlike dialanine, we perform an intermediate CG mapping where each backbone and sidechain of each amino acid is represented as separate CG sites (**Figures 3A and 4A**); note that glycine is mapped as a single backbone CG site. As a result, we obtain 111 and 51 LFT features for Trp-cage and chignolin, respectively.

**Figure 3.**
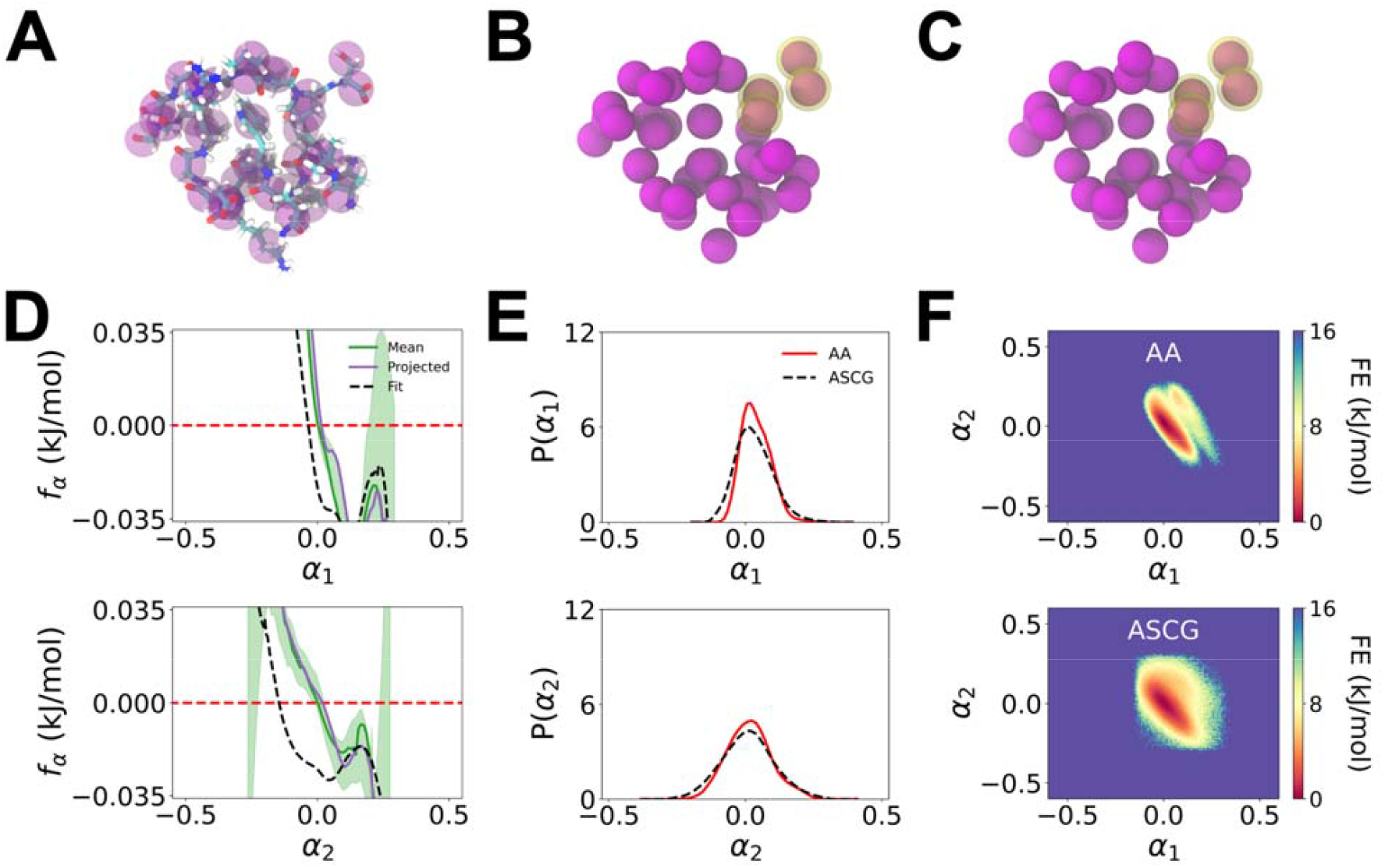
ASCG mapping, parameterization, and integration results for Trp-cage of the first two components. (A) Intermediate CG representation of Trp-cage using a center-of-mass mapping at a backbone-sidechain resolution, resulting in 37 CG sites shown in purple. (B-C) The ASCG mapping of (B) and (C) are depicted in yellow, where and both represent fluctuations localized to the C-terminus. (D) Force ( ) profiles for (top) and (bottom). The mean force represents the average forces along each ASCG component after transformation, shown with a 95% confidence interval. Projected forces represent forces estimated by calculating the 1D free energy surface from the 1D probability density of the atomistic data and then taking the negative gradient. Fit forces are predicted forces after fitting a multivariate nearest neighbors algorithm to the averaged ASCG transformed forces. (E) 1D probability distributions along (top) and (bottom), comparing the atomistic (AA) distribution with the ASCG distribution calculated after integrating and . (F) 2D slice of the free energy (FE) surfaces of and generated from AAMD trajectories (top) and ASCG trajectories after integration of and (bottom). The JSD between the underlying 2D probability distributions of and of the AAMD surface and the ASCG-generated surface is 0.075.

Our AAMD simulations show that Trp-cage maintains its folded structure over the entirety of the simulation at 270 K, with the root-mean-squared deviation (RMSD) of Trp-cage with respect to the experimental structure fluctuating between 1 to 4 Å over the course of the simulation (**Fig. S8B**). Our top seven ASCG components contain 34% explained variance. In **Figure 3**, we focus on the top two ASCG components, and the results for all seven ASCG components can be found in **Fig. S9**. Our top two ASCG components emphasize minimal movement outside of fluctuations observed near the C-terminus, as shown in **Figure 3B-C**. As shown for the top two ASCG components in **Figure 3D**, and for the top seven components in **Fig. S9B**, the mean effective ASCG forces for ASCG components are consistent with the negative gradient of their respective 1D component-wise PMFs. When integrating *α*_1_ through *α*_7_ over 100 independent replicas of 50 ns, we were able to achieve timesteps of 0.1 ps (and *γ* = 0.5 ps^-1^) without observing evidence of integration error. As AAMD simulations are typically limited to 1-2 fs, while conventional CGMD simulations are often limited to 10-25 fs, our ASCG variables demonstrate at least a factor of four to ten (two orders of magnitude) larger integration efficiency compared to that of CGMD (AAMD).

The Trp-cage configurations sampled by ASCG are consistent with those observed from AAMD. First, as seen in **Figure 3E**, the 1D ASCG distributions for and exhibit good agreement with that of the AAMD simulations with JSD values of 0.021 and 0.008, respectively. The 1D ASCG distributions of through also demonstrate excellent agreement (**Fig. S9B)**, with JSD values ranging from 0.003 to 0.051 for through . The 2D slices of the FESs depicted in **Fig. S9D** show that Trp-cage maintains one primary state (i.e., the folded state), with **Figure 3F** showing fluctuations in the directions of both and ; the metastable state at (0.0, -0.02) is representative of the fluctuations throughout the C-terminus, as this is the most flexible region in the atomistic Trp-cage simulations. The 2D JSD of and between the ASCG and AAMD distributions is 0.075, while the 7D JSD is 0.132. We note the higher JSD at higher dimensionality, which we attribute to the smoothing of the higher-dimensional FES. We observe that all 2D slices in **Fig. S9D** have low 2D JSD values, ranging from 0.033 to 0.089, indicating that the ASCG simulations faithfully recapitulate the AA conformational sampling.

Finally, we investigate ASCG for chignolin (**Figure 4A**), which undergoes rapid folding and unfolding at our simulated temperature of 420 K. The top five ASCG components for chignolin contain 34% explained variance, and we observe a drop in the spectral ratio after the fifth component. The corresponding mapping and forces of the first five ASCG components are shown in **Fig. S10A-B**, while we focus on the top two components in **Figures 4B-D**. Overall, we find that the top three ASCG components correspond to fluctuations around each of the two termini and find that *α*_4_ and *α*_5_ correspond to the central region of the protein. Together, these motions describe the transitions between the *β*-hairpin and coiled state of chignolin, with *α*_1_ representing the N-terminus side of the protein splaying away from and swinging back towards the C-terminus side during unfolding and folding, respectively, and *α*_2_ representing the equivalent splaying away and swinging in of the N-terminus during unfolding and folding, respectively. Unlike *α*_1_ and *α*_2_, *α*_3_ through *α*_5_ have a unimodal 1D probability distribution (**Fig. S10C**), and from the 2D slices of the FES (**Fig. S10D**), we note that the motions encoded by *α*_3_ in the C-terminus and by *α*_4_ and *α*_5_ in the central region of the protein are shared between the folded and unfolded states. While *α*_3_ through *α*_5_ components alone cannot distinguish between the two states, these components highlight motions of the protein that are mechanistically important to the transition between the two states, such as the side-chain interactions in the central region of chignolin that form turn-structures in chignolin.^81^ Similar to Trp-cage, we used a large timestep of *γ* = 0.1 ps with 0.5 ps^-1^ to integrate *α*_1_ through *α*_5_ for 100 independent replicas of 50 ns each. As seen in **Figures 4E-F** for the top two ASCG components, the ASCG model recapitulates both the 1D and 2D distributions from AAMD simulations with 1D JSDs of 0.007 and 0.005 for *α*_1_ and *α*_2_, respectively, and 2D JSD for *α*_1_ and *α*_2_ of 0.022. The 5D JSD for *α*_1_ through *α*_5_ is 0.037, showing excellent recapitulation even at higher dimensions. The 2D FESs depicted in **Figure 4F** show the folded state as a free energy minimum at ( *α*_1_, *α*_2_ ) = (0.14, - 0.24) while the unfolded state is the remaining broad region described by decreasing *α*_1_ and increasing *α*_2_. A similar confined folded state and broad region corresponding to the unfolded state is seen in 2D slices of the FES surface between other pairs of ASCG components in **Fig. S10D**. The time series profiles for *α*_1_ and *α*_2_ also exhibit reversible conformational switching between the folded and unfolded state that is consistent with the AAMD trajectories (**Fig. S11**).

**Figure 4.**
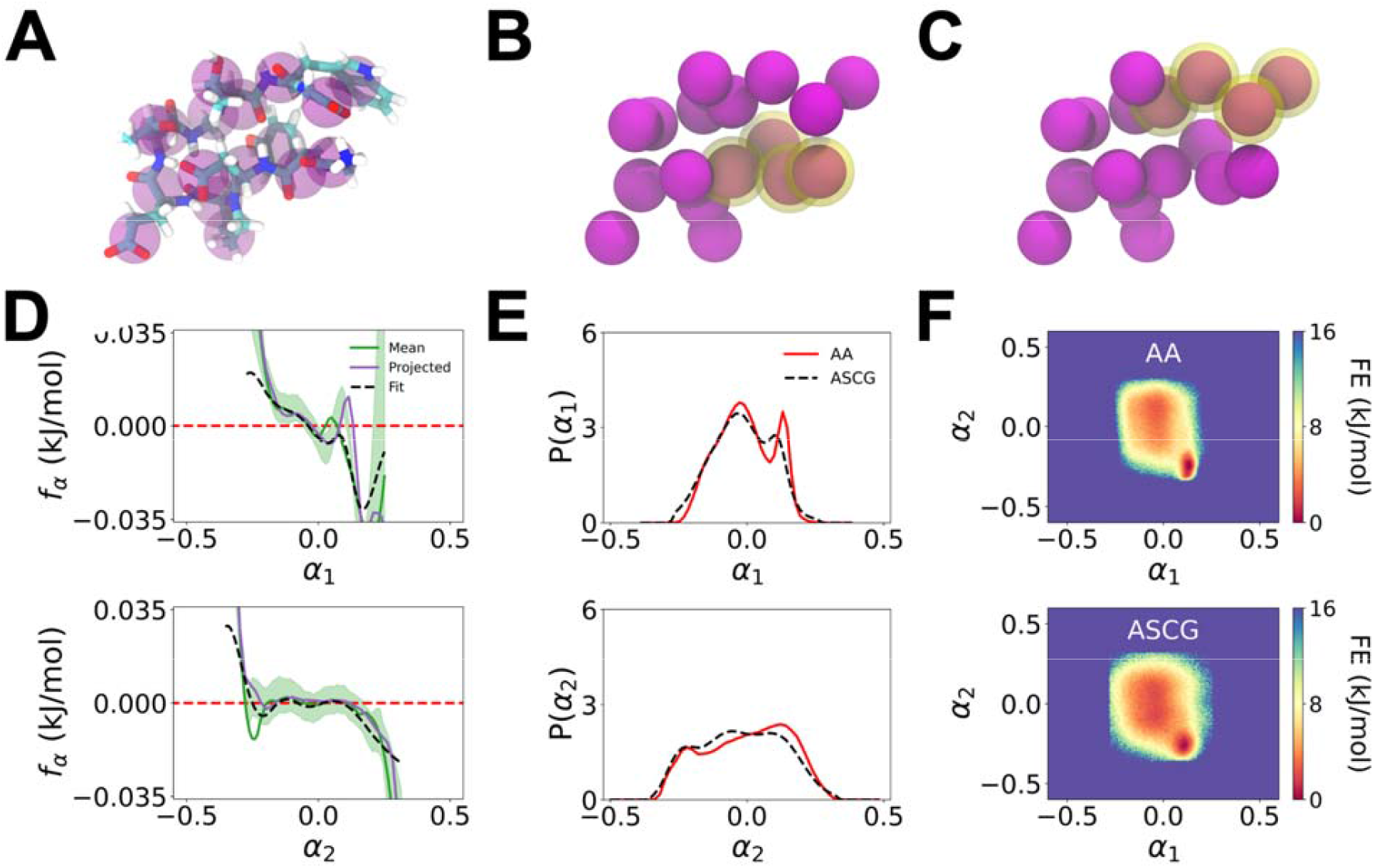
ASCG mapping, parameterization, and integration results for chignolin of the first two components. (A) Intermediate CG representation of chignolin using a center-of-mass mapping at a backbone-sidechain resolution, resulting in 17 CG sites shown in purple. (B-C) The ASCG mapping of (B) and (C) are depicted in yellow, where and represent the N-terminus and C-terminus, respectively. (D) Force ( ) profiles for (top) and (bottom). The mean force represents the average forces along each ASCG component after transformation, shown with a 95% confidence interval. Projected forces represent forces estimated by calculating the 1D free energy surface from the 1D probability density of the atomistic data and then taking the negative gradient. Fit forces are predicted forces after fitting a multivariate nearest neighbors algorithm to the averaged ASCG transformed forces. (E) 1D probability distributions along (top) and (bottom), comparing the atomistic (AA) distribution with the ASCG distribution calculated after integrating and . (F) 2D slice of the free energy (FE) surfaces of and generated from AAMD trajectories (top) and ASCG trajectories after integration of *α*_1_ and *α*_2_ (bottom). The JSD between the underlying 2D probability distributions of *α* _1_ and *α*_2_ of the AAMD surface and the ASCG-generated surface is 0.022.

In summary, the ASCG framework is able to both identify and simulate the major conformational motions of the miniproteins Trp-cage and chignolin. In the former, the conformational motions of Trp-cage are largely suppressed at 270 K and ASCG identifies only local fluctuations around one major (i.e., folded) conformational state. In the latter, ASCG identifies the reversible folding and unfolding of chignolin at an elevated temperature of 420 K.

#### Robustness to Limited Training Data

Next, we tested whether the ASCG framework is robust to limited AAMD data provided for training, such as when computational resources are limited. To test the minimum amount of data needed, we repeated the ASCG workflow using shorter AAMD simulation times for training. We measured the performance with two types of JSD values: the ASCG distribution compared against the distribution from the full (untruncated) reference AAMD dataset and the distribution from the shortened AAMD trajectories used in training. Before measuring JSD of the distribution of ASCG models trained with truncated datasets against the distribution of the full reference AAMD dataset, we checked for whether a rotation of the truncated FES was needed, as mentioned in *Theory and Methods*.

We find that the ASCG workflow excels at recapitulating the distribution of the training data from which it was trained, reflected in **Figure 5** by the low JSDs of the purple training curves. If the distribution of the shortened AAMD trajectory is consistent with that of the full AAMD dataset, then we also expect the ASCG models trained from truncated data to yield low JSDs when compared to the full AAMD statistics.

**Figure 5.**
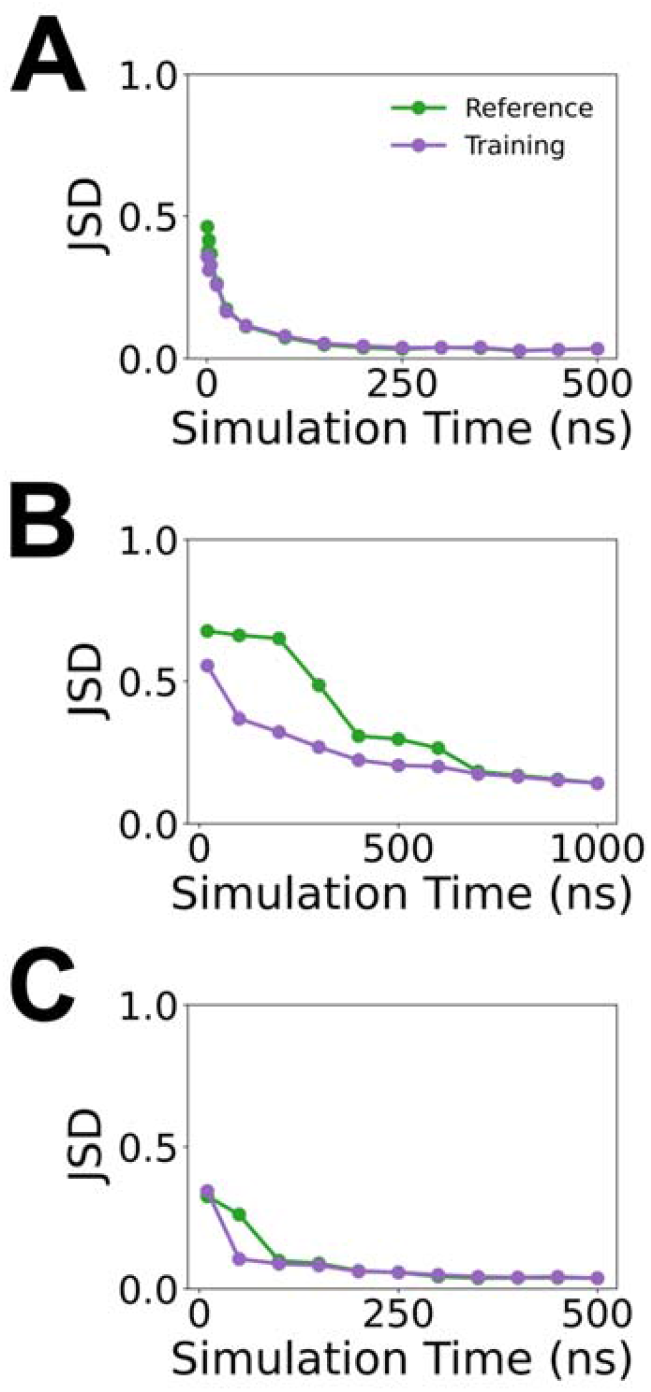
Effect of reduced training data on ASCG performance. (A-C) The JSD of the underlying probability distributions as a function of total AAMD simulation time used for training of (A) dialanine, (B) Trp-cage, and (C) chignolin models. The JSD is calculated between the 3D, 7D, and 5D distributions for dialanine, Trp-cage, and chignolin, respectively, from the ASCG-generated trajectories in comparison to the full reference AAMD dataset (green) and the truncated AAMD dataset that was used to train ASCG (purple).

For dialanine (**Figure 5A**), a minimum of 150 ns of AAMD data appears to be necessary to quantitatively reproduce the distribution of the truncated dataset (with a 3D JSD of 0.053) and qualitatively, the full 500 ns dataset (with a 3D JSD of 0.049). While low 3D JSD values are measured when using as little as 25 ns, we start to observe sampling of unphysical space. Below 25 ns, the truncated dataset distribution diverges from that of the full-length AAMD dataset (**Fig. S12**). For Trp-cage (**Figure 5B**), the ASCG method resulted in excellent agreement with the full 1,000 ns AAMD trajectory until down to 400 ns of training data. At shorter simulation times, the truncated AAMD distribution diverges from that of the full AAMD distribution (**Fig. S13**) and thus results in a large increase of the JSD with respect to the full dataset. For chignolin (**Figure 5C**), when limiting the ASCG training to just the first 100 ns of data, the 5D JSD is 0.086 with respect to the training dataset and 0.099 with respect to the full 500 ns dataset. Using the first 50 ns of data, the 5D JSD is 0.105 with respect to the truncated dataset, although we see start to see the impacts of limited training data as the 5D JSD more than doubles to 0.261 with respect to the full dataset. Thus, we treat 100 ns as the data reduction limit for chignolin. Overall, we note that the ASCG method recapitulates the training data and is still able to sample the transitions between the folded and unfolded states even when training data is reduced by an order of magnitude (**Fig. S14**).

In summary, the ASCG framework is successfully able to recapitulate distributions even when training data is limited but is unable to extrapolate beyond the training data itself. We found that for dialanine and chignolin, the training data could be shortened by an order of magnitude to 150 ns and 100 ns, respectively. However, at least 400 ns of data was still required for Trp-cage; we speculate that the larger data requirements for Trp-cage are a result of ASCG discovering a subtle and slow fluctuation near the C-terminus, as we expect Trp-cage to be largely stationary.

#### Comparison of ASCG to PCA

The ASCG workflow bears some similarity to PCA, except the former is a supervised dimension reduction technique while the latter is unsupervised. As such, we next compared ASCG to PCA, where we specifically tested whether PCA eigenvectors could also be used as a linear operator for both mapping and effective forces, using the same number of PCA components as used for ASCG for each system (three, seven, and five for dialanine, Trp-cage, and chignolin, respectively). We used the same hyperparameters for nearest neighbor multivariate force interpolation with PCA as for ASCG, choosing a bin width that results in a similar number of samples that was used in the ASCG force interpolation, to ensure similar complexity of interpolation schemes for PCA and ASCG frameworks. This resulted in a bin width of 0.1, 0.2, and 0.15 for dialanine, chignolin, and Trp-cage, respectively. For chignolin, we additionally tested whether PCA is robust to limited data like our proposed ASCG framework.

We find that for dialanine (**Fig. S15**) and Trp-cage (**Fig. S16**), PCA can recapitulate atomistic distributions, suggesting that variance in positions and variance in forces are coupled in these cases. However, notable differences between the ASCG and PCA models emerge in the case of chignolin (**Figure 6 and Fig. S17**), which is also the system with the most significant conformational changes amongst the three. Hence, for the remainder of this section, we will focus our discussion on chignolin.

**Figure 6.**
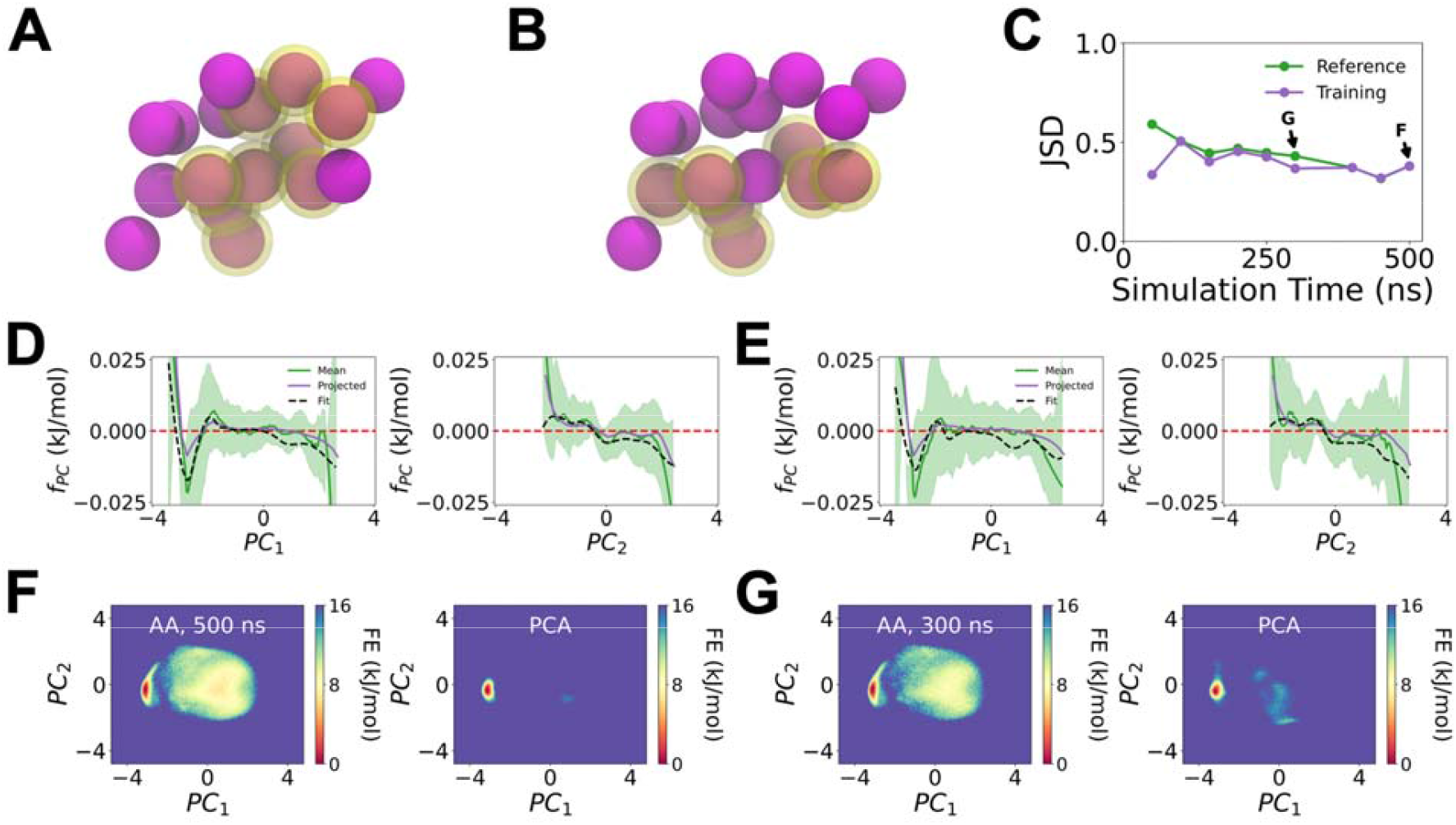
PCA as an alternative unified coarse-graining framework for chignolin, shown for the first two components. (A-B) Mapping determined by (A) and (B) on chignolin that was first coarse-grained at a backbone-sidechain resolution, where represented CG sites for each PC are highlighted in yellow. (C) The JSD as a function of AAMD simulation time used to train PCA models. The JSDs between the 5D probability distributions of the PCA-generated trajectories is plotted in comparison to the full reference dataset (green) and the shortened dataset that was used to train PCA (purple). (D-E) Force ( profiles along (left) and (right) when PCA was fit on (D) the full 500 ns of AAMD data and (E) a reduced simulation time of 300 ns. The mean force represents the average forces along each PC component after transformation, shown with a 95% confidence interval. Projected forces represent forces estimated by calculating the 1D free energy surface from the 1D probability density of the atomistic data and then taking the negative gradient. Fit forces are predicted forces after fitting a multivariate nearest neighbors algorithm to the averaged PCA-transformed forces. (F-G) The 2D free energy (FE) surfaces of and generated from AAMD trajectories (left) and from PCA trajectories after integration of and (right) using training data of length (F) 500 ns and (G) 300 ns. The JSD between the underlying 2D probability distributions of the full-length AAMD trajectory and the PCA-generated trajectory is 0.193 and 0.148 for 500 ns and 300 ns, respectively.

We find that the top five PCs for chignolin have 37% explained variance. As shown in **Figures 6A-B**, the top PC maps to a motion involving both sides of chignolin while the second PC maps primarily to the N-terminal half of chignolin. When using the full 500 ns AAMD dataset, the resulting PCA model yields a 2D JSD between PC1 and PC2 of 0.173 and 5D JSD of 0.381. We primarily see the PCA model sample the folded state (**Figures 6C and 6F**) with limited exploration of the unfolded state, which we also find in the 2D slices between remaining pairs of PCs (**Fig. S17D**). Also, as seen in **Figure 6C**, we observe that the performance of the PCA model generally worsens as the AAMD training data is truncated, although performance was already poor with the full-length dataset (refer to **Figure 5C**). We attribute the poor performance to the differences between the PCA-determined forces and the negative gradient of their respective 1D component-wise PMFs, such as the magnitude of the mean effective PC1 force being overestimated at the location of the minima (-2.80), seen in both the PC1 force profiles trained using 500 ns (**Figure 6D**) and 300 ns (**Figure 6E**) of data, leading to overstabilization of the folded state. In comparing the force profiles using 500 ns and 300 ns of data, we also see that limiting data has the effect of smoothing out the PCA-projected forces, which is most evident in PC1. With the full 500 ns of data, the PCA model primarily samples the minima at ( *PC*_1_, *PC*_2_) = (-2.80, -0.34), representing the folded region, with very limited exploration of the broad region in the FES, mainly clustering around the highest probable configuration in the unfolded state (**Figure 6F**). However, as seen in **Figure 6G**, we see that, in comparison to the model trained with 500 ns of data, the smoothing of the force profile using 300 ns of training data starts to result in unphysical sampling behavior in the PCA-generated trajectories, overestimation of the free energy barrier between the folded and unfolded states, and more sampling of the unfolded region. Hence, while the 2D FES using 300 ns of data is consistent with that of the 500 ns dataset, the trained PCA model is unable to faithfully recapitulate the AAMD distribution, with a measured 5D JSD of 0.368.

In summary, while PCA may be a suitable alternative to ASCG for simple systems like dialanine or static proteins like Trp-cage, PCA appears to be limited in its application for systems with large conformational changes or disordered regions, particularly when AAMD data may be limited. We attribute the success of ASCG over PCA in the case of chignolin to the supervised aspect of the ASCG method, which identifies collective motions that contribute to the greatest variance in the PMF rather than collective motions with the largest fluctuations.

## Conclusions

We demonstrated the use of active subspace learning for bottom-up coarse-graining as a unified framework for CG mapping and parameterization. Our proposed ASCG (or active subspace for coarse-graining) workflow identifies linear projections of configurational degrees of freedom that maximally describe the gradients of the potential energy, as sampled from AAMD simulations. The linear projection is an interpretable CG mapping operator for both CG variables and associated effective interactions (via force mapping). In other words, ASCG removes the need for separate *ad hoc* decisions on CG mapping and parameterization strategies and, by design, emphasizes collective motions that describe large changes to the underlying potential and free energy surfaces.

The ASCG method offers several advantages in computational efficiency compared to traditional, particle-based CG methods. The first benefit is that the ASCG methods dramatically reduce the dimensionality of the underlying model as each ASCG variable encodes collective motions rather than individual particles that participate in each collective motion. Assuming a CG resolution of one CG site per residue (which is considered a low-resolution model), the ASCG models for chignolin and Trp-cage with five and seven ASCG degrees of freedom, respectively, represent ∼17% and ∼12% of the typical CG degrees of freedom (30 and 60, respectively, after solvent is removed). The second benefit is that the integration timestep accessible to the ASCG model (∼100 fs) is a factor of four to ten times larger than that of traditional CG simulations, which tend to use 10-25 fs timesteps.^22, 25^ Finally, the third benefit is that ASCG model training is robust to limited training data. However, one limitation of the ASCG model is that it cannot extrapolate beyond the phase space sampled within the training data, which is also true of many other bottom-up CG methods. We expect that enhanced sampling methods combined with AAMD can bridge this limitation when combined with reweighting techniques.^82-84^

The current ASCG workflow has multiple opportunities for refinement and future development. We are currently using a multivariate force interpolator that implements *N*-dimensional binning, where *N* is the number of ASCG components. For larger systems with complex motions that require an increased number of ASCG components, limitations due to the curse of dimensionality would become evident, where the training becomes increasingly sparse due to the increase in volume of higher-dimensional space. The limitation of sparse input data started to be evident in Trp-cage when using the top seven ASCG components, where we observed a loss of detail in the ASCG FES in comparison to the reference FES from AAMD. To address this, alternative multivariate force models should be explored, e.g., an artificial neural network. Additionally, the LFT-transformed Cartesian positions we used as features may not be expressive enough for a clear spectral gap to be evident in the case of larger biomolecules. To this end, we are currently investigating other input basis functions, both to represent intramolecular interactions within larger biomolecules and to capture intermolecular interactions between biomolecules, as well as nonlinear extensions to ASCG based on the kernel trick^85^. Finally, one aspect of coarse-graining that we have yet to discuss is backmapping. The recovery of high-dimensional atomistic details from low-dimensional CG simulations, for example to determine specific residue-residue contacts, is inherently a difficult task.^86^ However, data-driven approaches have recently shown promise for backmapping from traditional CG models^87-90^ and we anticipate that similar methods can be applied for ASCG-to-AA backmapping.

In summary, the ASCG method is a step forward toward a unified bottom-up coarse-graining framework. Rather than coarse-graining new pseudo-particles as done in traditional CG methods, we leverage active subspace learning to identify collective motions representing the largest changes to the underlying potential (and free) energy surface, as well as associated forces and equations of motion describing changes to these collective motions. In the future, we anticipate that the ASCG method will facilitate long timescale simulations of biological and soft matter systems that are currently inaccessible to both AA and CGMD simulations.

## Supplemental Materials

Additional details on featurization, analysis of AAMD trajectories, and characterization of alternative ASCG and PCA models can be found online.

## Acknowledgements

This work was partially supported by the National Institutes of Health through grant R35GM157192 (for A.W. and A.J.P.) and by the National Science Foundation through grant DMS-2107938 (for S.P.). This work used high-performance computing resources made available by the Research Computing Group at Colorado School of Mines and from Anvil at the Rosen Center for Advanced Computing through allocation BIO220015 from the Advanced Cyberinfrastructure Coordination Ecosystem: Services & Support (ACCESS) program, which is supported by National Science Foundation grants #2138259, #2138286, #2138307, #2137603, and #2138296.

## Data Availability Statement

The data underlying this study, including simulation files, analysis scripts, and processed data, are openly available on GitLab: https://gitlab.com/pak-group/ascg_m01.

