## Supplementary Information for "Active subspace learning for coarse-grained molecular dynamics"

### Maintaining roto-translational invariance using a local frame transformation

We required the coordinates and forces used to train our ASCG models to be translationally and rotationally invariant. To do this, we adapted an approach used by DeePCG called the local frame transformation (LFT)[1].

We apply the LFT to each frame of the unscaled atomistic data prior to applying mass weighting or centering the features. For each system, we take three reference sites, consecutive CG sites which we label with indices $i$, $j$, $k$. To enforce translational invariance, we first translate the coordinates of each frame such that reference site $j$ is the origin. Forces are inherently translationally invariant, so we do not apply this translation to the CG cartesian forces. The entire frame is then rotated to align the $xy$ plane to the plane defined by CG sites $i$, $j$, $k$. The transformation is as follows:

$$x = R\mathrm{X} = \begin{bmatrix} \overrightarrow{r_{jk}} \\ \overrightarrow{r_{jk}} \times (\overrightarrow{r_{jk}} \times \overrightarrow{r_{ji}}) \\ (\overrightarrow{r_{jk}} \times \overrightarrow{r_{ji}}) \end{bmatrix} \begin{bmatrix} \mathrm{x}_1 & \dots & \mathrm{x}_\mathrm{n} \\ \mathrm{y}_1 & \dots & \mathrm{y}_\mathrm{n} \\ \mathrm{z}_1 & \dots & \mathrm{z}_\mathrm{n} \end{bmatrix} \tag{1}$$

$$f_x = R\mathrm{F}_\mathrm{X} = \begin{bmatrix} \overrightarrow{r_{jk}} \\ \overrightarrow{r_{jk}} \times (\overrightarrow{r_{jk}} \times \overrightarrow{r_{ji}}) \\ (\overrightarrow{r_{jk}} \times \overrightarrow{r_{ji}}) \end{bmatrix} \begin{bmatrix} \mathrm{f}_{\mathrm{x}_1} & \dots & \mathrm{f}_{\mathrm{x}_\mathrm{n}} \\ \mathrm{f}_{\mathrm{y}_1} & \dots & \mathrm{f}_{\mathrm{y}_\mathrm{n}} \\ \mathrm{f}_{\mathrm{z}_1} & \dots & \mathrm{f}_{\mathrm{z}_\mathrm{n}} \end{bmatrix} \tag{2}$$

where $\boldsymbol{x}$ is the matrix of LFT CG site coordinates, $\boldsymbol{f_x}$ is the corresponding LFT matrix of CG site forces, $R$ is the rotation matrix defined by the vectors $\overrightarrow{\boldsymbol{r_{jk}}}$ and $\overrightarrow{\boldsymbol{r_{ji}}}$ between the CG reference atom coordinates, $X$ is the matrix of translated cartesian CG site coordinates, and $F_X$ is the matrix of corresponding cartesian CG forces. We note that when translating the system to the origin and applying the transformation, the total degrees of freedom in each system is reduced by 6, i.e., 6 of the features are always 0.0.

We also note that the selection of reference sites may change the motions that are identified by the ASCG framework. When aligning the system to different reference sites, we noticed that poorly selected sets of reference sites resulted in atoms flipping around an axis within one frame, leading to an artificially large change in the LFT coordinates. To address this, we swept through all combinations of possible reference sites for each system, selecting the set of reference sites with the lowest means of LFT feature variances.

**Correcting rotation of truncated dataset probability densities**

When testing the critical point of the ASCG framework to limited training data, we qualitatively observed a rotation of the ND FES in truncated training datasets, most notably observed for dialanine (**Fig. S1**). We attributed the large increase in JSD between ASCG-generated and the full-length AAMD distributions as being due to the rotation of the ASCG phase space upon training from shortened trajectories. To correct the rotation, we treated the ND eigenvector matrix of the full-length trajectory and truncated trajectory as an orthogonal Procrustes problem[2].

To goal of the orthogonal Procrustes problem is to find an orthogonal matrix that most closely matches a given matrix into another given matrix with an orthogonal transformation matrix[2]. We aim to transform the active subspace from the truncated trajectory (denoted by $W_{1t} \in \mathbb{R}^{3n \times m}$) to the active subspace matrix determined from the full-length trajectory (denoted by $W_{1f} \in \mathbb{R}^{3n \times m}$). First, we let $R \in \mathbb{R}^{m \times m}$ denote the rotation matrix that most closely matches $W_{1t}$ to $W_{1f}$, under the condition that $R^T R = I$. Under this condition for R, we aim to minimize the sum of squares residual matrix[2]:

$$E = W_{1t}R - W_{1f} \tag{3}$$

We state the minimization problem as:

$$tr(E^T E) = min \tag{4}$$

This can be formulated as follows[2]:

$$tr(E^T E) = tr(\left(W_{1t}R - W_{1f}\right)^T (W_{1t}R - W_{1f})) \tag{5}$$

$$tr(E^T E) = tr(R^T W_{1t}^T W_{1t} R) + tr\left(W_{1f}^T W_{1f}\right) - 2tr\left(R^T W_{1t}^T W_{1f}\right) \tag{6}$$

Where $tr(R^T W_{1t}^T W_{1t} R) + \ tr\left(W_{1f}^T W_{1t}^T\right)$ in Eq. 7 reduces to a constant and we will denote as $K$.

$$tr(E^T E) = K - 2tr\left(R^T W_{1t}^T W_{1f}\right) \tag{7}$$

As shown in Eq. 8, we now solve for the maximum of $2tr\left(R^T W_{1t}^T W_{1f}\right)$. To do this, we find the spectral decomposition of $W_{1t}^T W_{1f}$.

$$W_{1t}^T W_{1f} = \ U \Lambda V^T \tag{8}$$

From which we construct the rotation matrix $R = UV^T$ and we apply the rotation directly to the ASCG trajectories generated from truncated datasets.

$$\widehat{\alpha_{ASCG}} = \ \alpha_{ASCG} R \tag{9}$$

The resulting rotated ASCG trajectory, denoted by $\widehat{\alpha_{ASCG}}$, is then used to calculate the ND JSD between the full-length AAMD trajectories for analyzing the effect of limited training data on ASCG performance. If an improvement in the JSD is seen after applying the proposed rotation matrix, the rotation is accepted, and the JSD measured after applying a rotation is recorded. If the proposed rotation does not improve or worsen the JSD, the rotation is rejected and the JSD measured before the rotation is kept. An example comparing FES when using the original or rotated ASCG trajectories for dialanine can be found in **Fig. S1**, as well as a comparison of the JSD calculated using the original or rotated ASCG trajectory between the full-length atomistic trajectory.

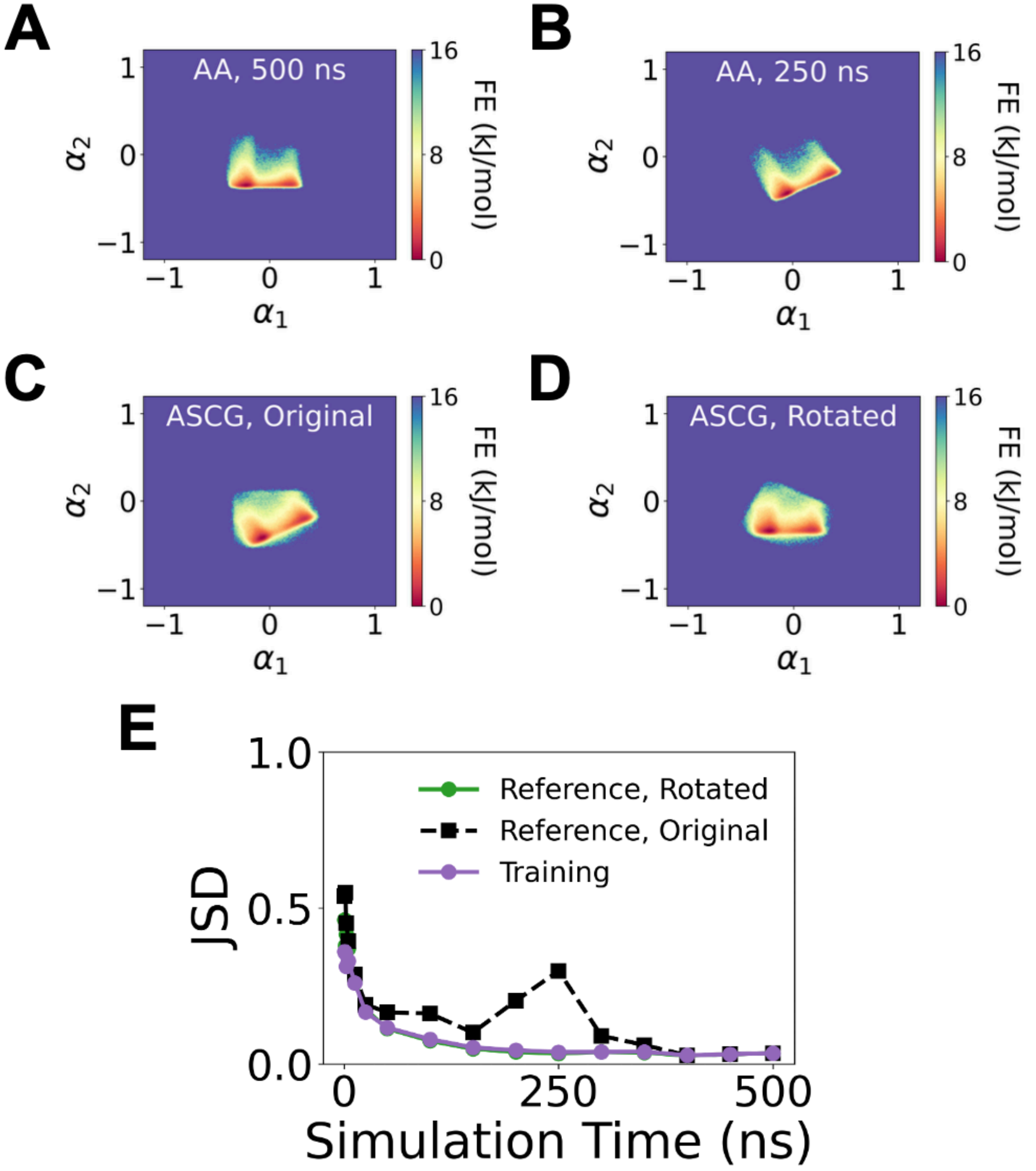


**Fig. S1.** Rotation of the dialanine distribution when the simulation time is shortened. The full effective simulation time of dialanine is 500 ns, showing two minima in the A) atomistic 2D free energy surface. Qualitatively, the two minima of the first 250 ns in the B) 2D atomistic free energy surface look to be a rotation of (A). C) shows excellent recapitulation in the ASCG-generated 2D free energy surface to (B). After rotating the ASCG-generated free energy surface according to the rotation matrix determined by the orthogonal Procrustes solution, the 2D free energy surface has excellent recapitulation to (A). D) The JSD as a function of total AAMD simulation time used for training of the dialanine model. The JSD is calculated between the 3D distributions from the ASCG-generated trajectories in comparison to the full reference AAMD dataset before applying the rotation matrix (black) and after applying the rotation matrix (green), and the truncated AAMD dataset (purple).

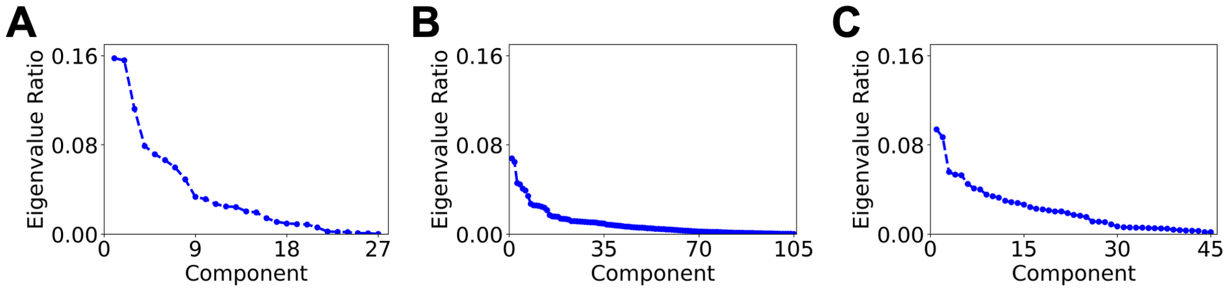


**Fig. S2.** Eigenvalue ratio spectra for A) dialanine, B) Trp-cage, and C) chignolin obtained from spectral decomposition of the normalized covariance matrix of effective forces $\hat{\boldsymbol{f}}$ along $\hat{\boldsymbol{x}}$ .

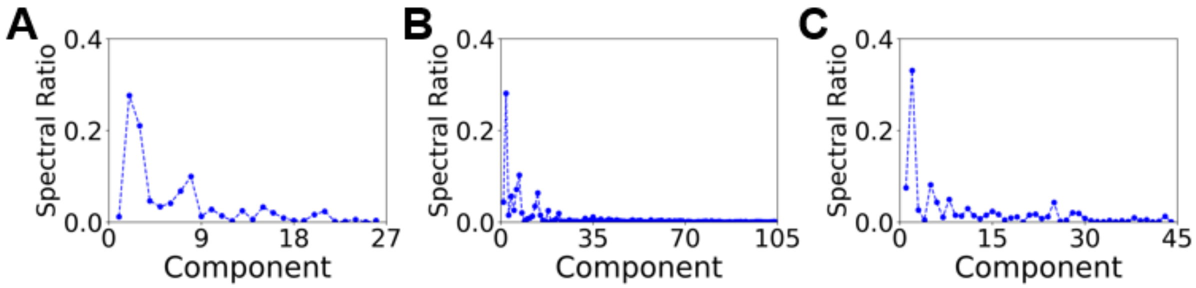


**Fig S3.** Spectral ratio of eigenvalues $(\Lambda_i - \Lambda_{i+1})/\Lambda_1$ after each ASCG component for A) dialanine, B) Trp-cage, and C) chignolin.

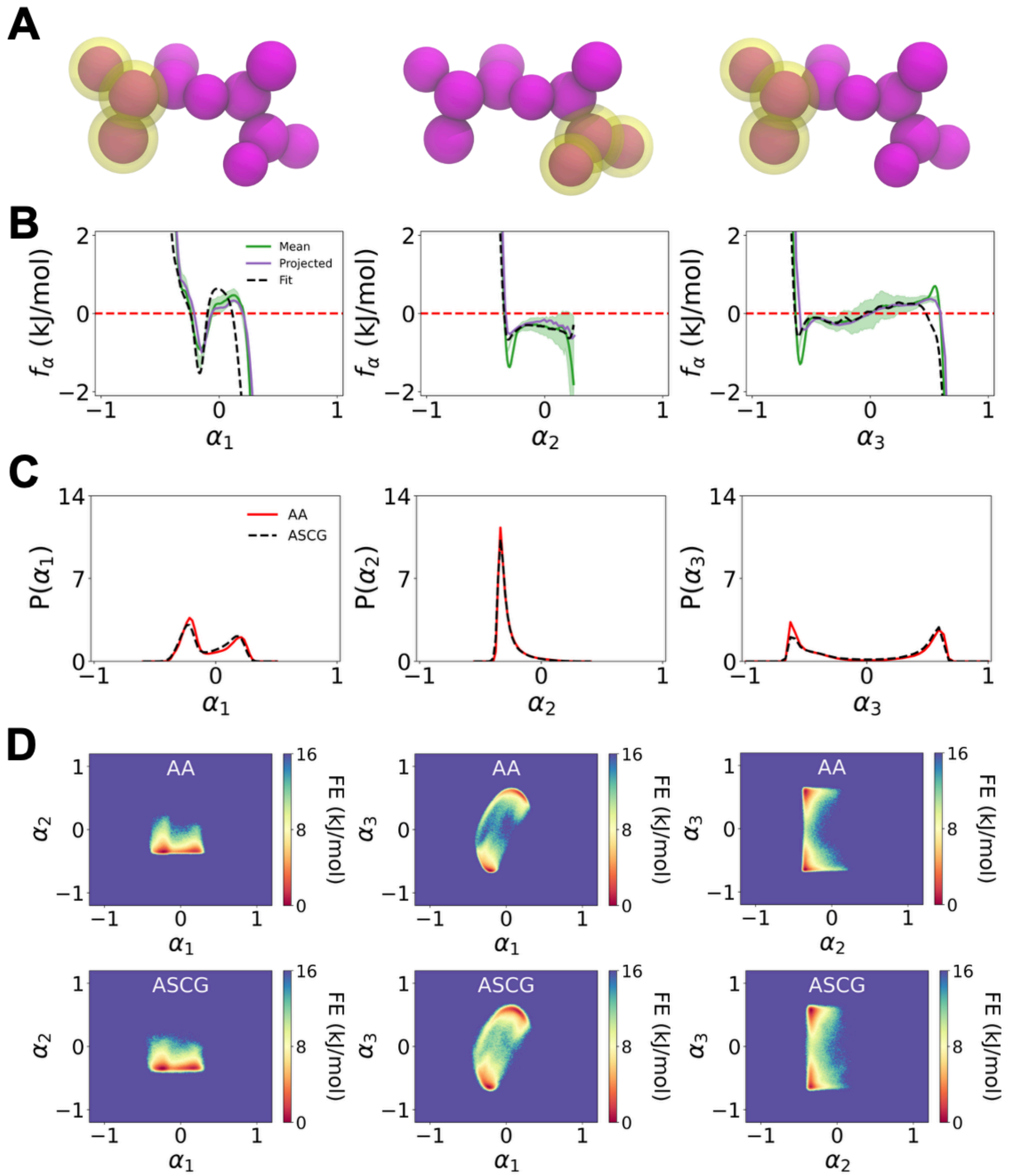


**Fig. S4.** Complete mapping, parametrization, and integration results of 3 ASCG components for dialanine. (A) ASCG mapping of $\alpha_1$ (left), $\alpha_2$ (middle), and $\alpha_3$ (right) depicted in yellow with the 11 united atom resolution CG sites shown in purple. (B) Force ($f_\alpha$) profiles for $\alpha_1$ (left), $\alpha_2$ (middle), and $\alpha_3$ (right). The mean force represents the average forces along each ASCG component after transformation, shown with a 95% confidence interval. Projected forces represent forces estimated by calculating a 1D free energy surface from the 1D probability density of the atomistic data and then taking the negative gradient. Fit forces are predicted forces after fitting a multivariate nearest neighbors algorithm to the averaged ASCG transformed forces. (C) 1D probability distributions along $\alpha_1$ (left), $\alpha_2$ (middle), and $\alpha_3$ (right) comparing the atomistic (AA) distribution with the ASCG distribution calculated after integrating $\alpha_1$, $\alpha_2$, and $\alpha_3$. (D) 2D slices of the 3D free energy (FE) surface of $\alpha_1$ and $\alpha_2$ (left), $\alpha_1$ and $\alpha_3$ (middle), and $\alpha_2$ and $\alpha_3$ (right), generated from AAMD trajectories (top) and from ASCG integration (bottom). The JSD between the underlying 3D probability distributions of the AAMD surface and the ASCG-generated surface is 0.034.

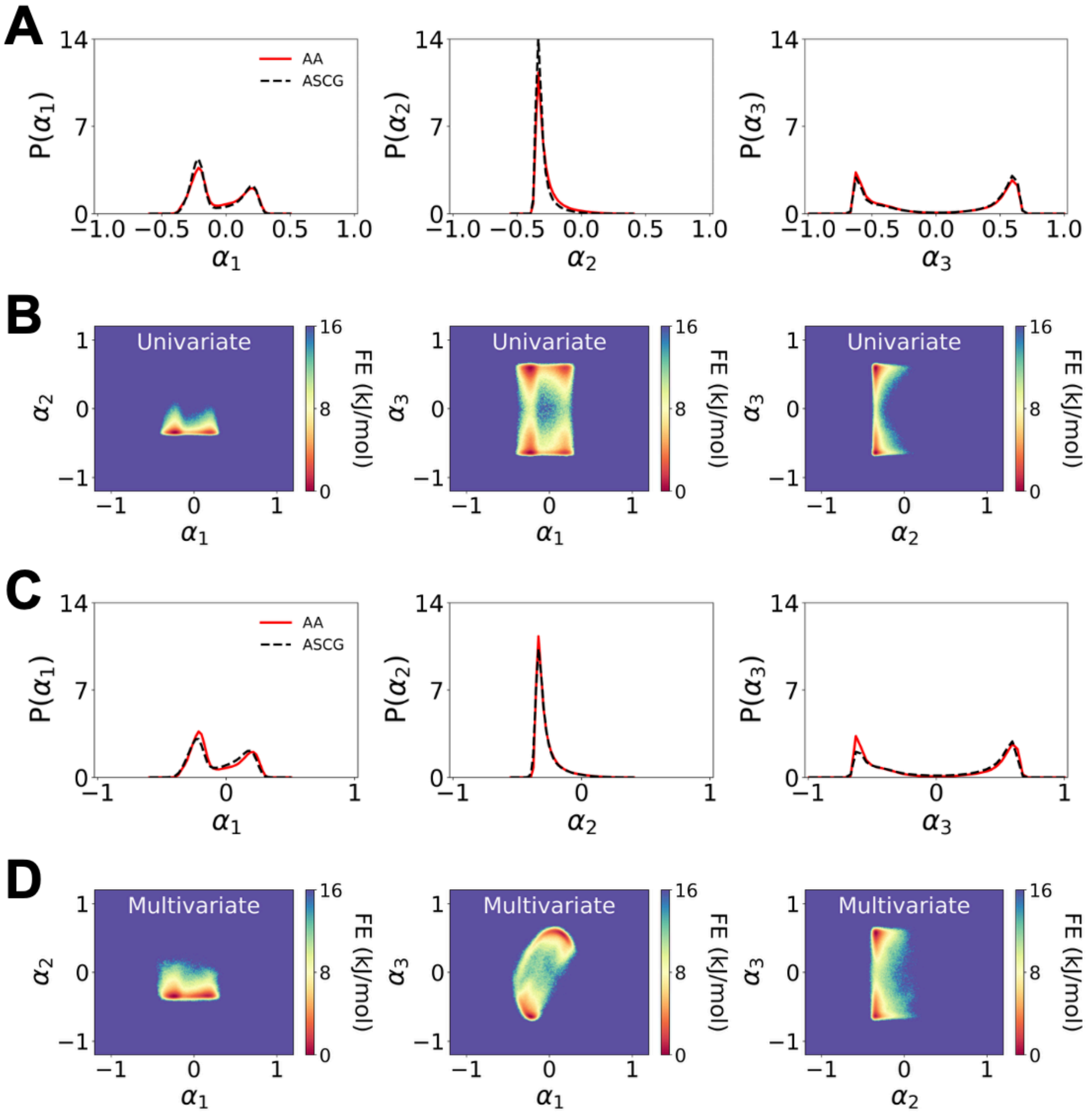


**Fig. S5** Comparison of univariate and multivariate force interpolation schemes for dialanine. Univariate: A) 1D probability distribution for $\alpha_1$ (left), $\alpha_2$ (middle) and $\alpha_3$ (right), and B) 2D slices of the 3D free energy surface (FES) generated from ASCG integrated trajectories using a univariate force algorithm. The underlying probability distribution of the 3D FES has a JSD of 0.196 with respect to the atomistic probability distribution. Multivariate: A) 1D probability distribution for $\alpha_1$ (left), $\alpha_2$ (middle) and $\alpha_3$ (right), and B) 2D slices of the 3D FES generated from ASCG integrated trajectories using a multivariate force algorithm. The underlying probability distribution of the 3D FES has a JSD of 0.034 with respect to the atomistic probability distribution.

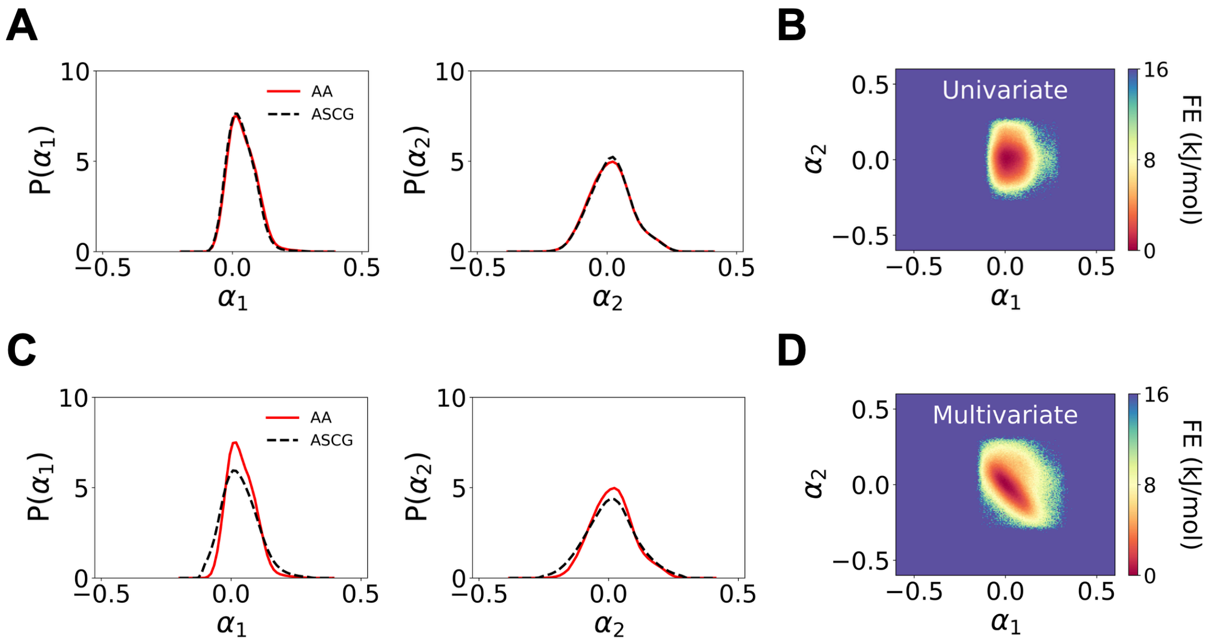


**Fig. S6.** Comparison of univariate and multivariate force interpolation schemes for Trp-cage, focusing on the first two ASCG components. Univariate: A) 1D probability distribution for $\alpha_1$ (left) and $\alpha_2$ (right), and B) 2D free energy surface generated from ASCG integrated trajectories using a univariate force algorithm. The underlying probability distribution of the 7D FES has a JSD of 0.209 with respect to the atomistic probability distribution. Multivariate: A) 1D probability distribution for $\alpha_1$ (left) and $\alpha_2$ (right), and B) 2D slice of the FES of $\alpha_1$ and $\alpha_2$ generated from ASCG integrated trajectories using a multivariate force algorithm. The underlying probability distribution of the 7D FES has a JSD of 0.132 with respect to the atomistic probability distribution.

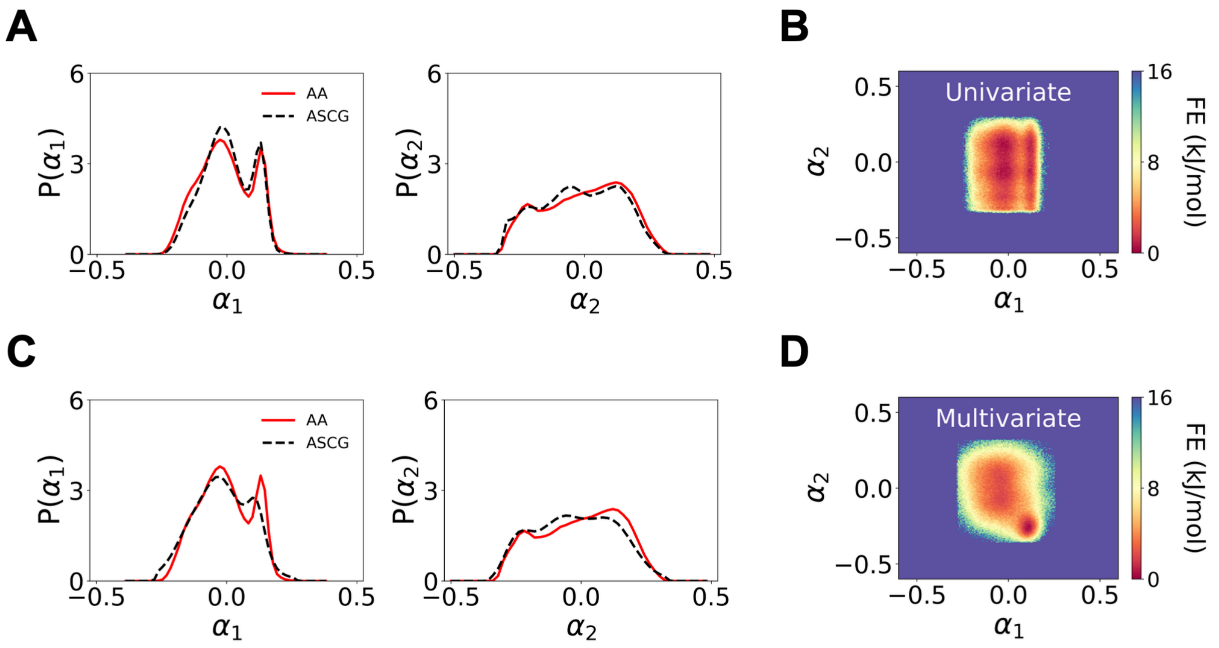


**Fig. S7.** Comparison of univariate and multivariate force interpolation schemes for chignolin, focusing on the first two ASCG components. Univariate: A) 1D probability distribution for $\alpha_1$ (left) and $\alpha_2$ (right), and B) 2D slice of the free energy surface generated from ASCG integrated trajectories using a univariate force algorithm. The underlying probability distribution of the 5D FES has a JSD of 0.095 with respect to the atomistic probability distribution. Multivariate A) 1D probability distribution for $\alpha_1$ (left) and $\alpha_2$ (right), and B) 2D slice of the original 5D FES of $\alpha_1$ and $\alpha_2$ generated from ASCG integrated trajectories using a multivariate force algorithm. The underlying probability distribution of the 5D FES has a JSD of 0.037 with respect to the atomistic probability distribution.

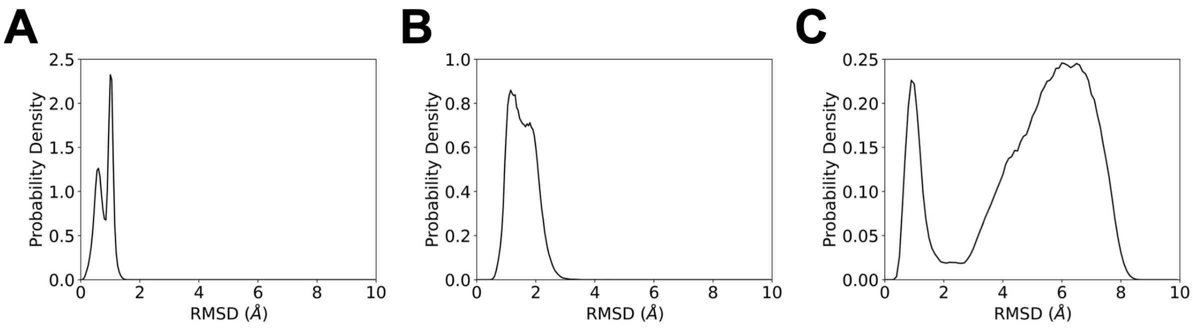


**Fig. S8.** Probability density of the root-mean-squared-displacement (RMSD) with respect to experimental structures across all AAMD data for A) dialanine, B) Trp-cage, and C) chignolin.

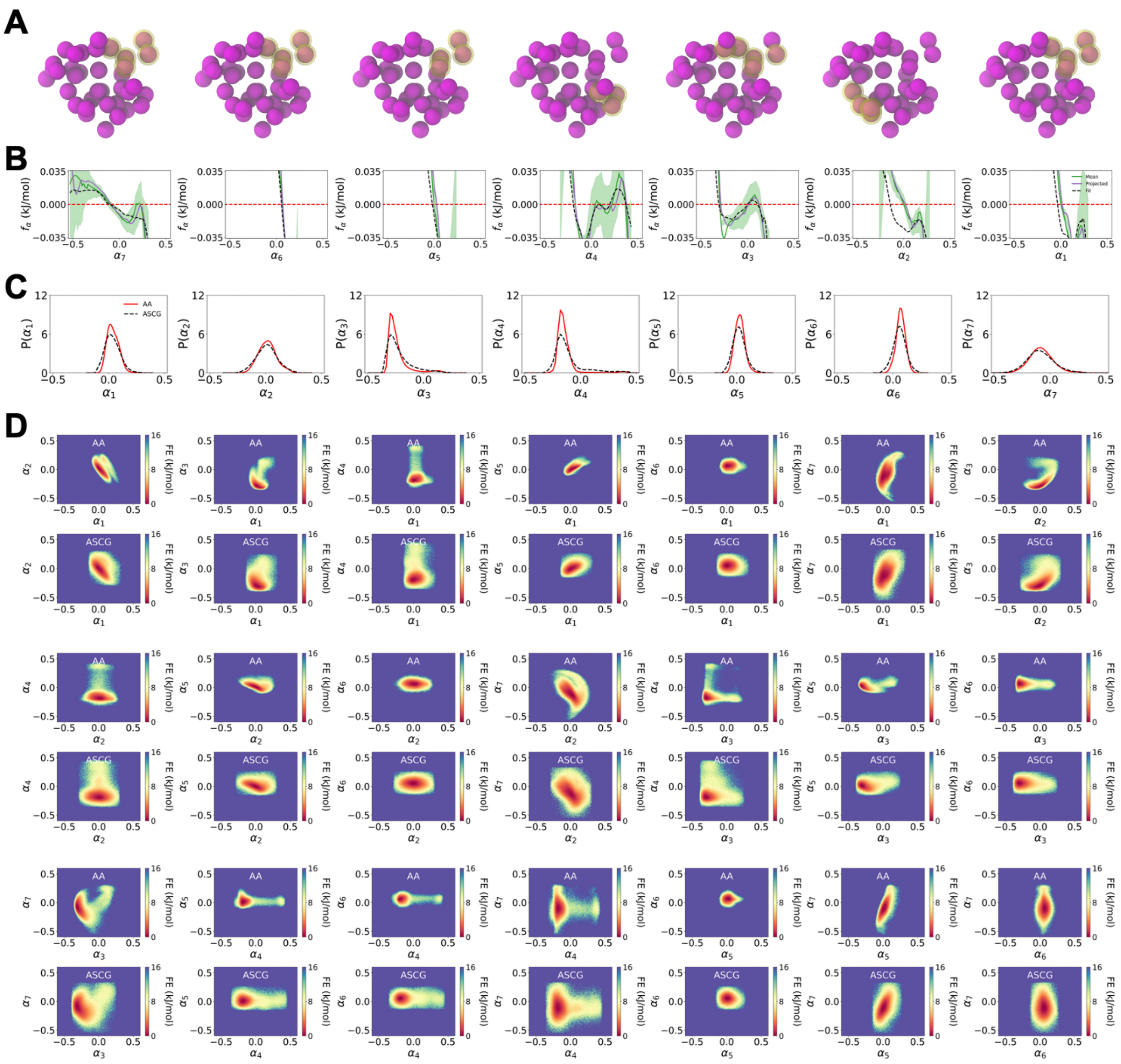


**Fig. S9.** Complete mapping, parametrization, and integration results of 7 ASCG components for trp-cage. (A) ASCG mapping of $\alpha_1$ through $\alpha_7$, in order from left to right, depicted in yellow. (B) Force ($f_\alpha$) profiles for $\alpha_1$ through $\alpha_7$. The mean force represents the average forces along each ASCG component after transformation, shown with a 95% confidence interval. Projected forces represent forces estimated by calculating a 1D free energy surface from the 1D probability density of the atomistic data and then taking the negative gradient. Fit forces are predicted forces after fitting a multivariate nearest neighbors algorithm to the averaged ASCG transformed forces. (C) 1D probability distributions along $\alpha_1$ through $\alpha_7$ comparing the atomistic (AA) distribution with the ASCG distribution calculated after integrating $\alpha_1$ through $\alpha_7$. (D) 2D slices of the 7D free energy (FE) surface between each pair of ASCG components, generated from AAMD trajectories (top) and from ASCG integration (bottom). The JSD between the underlying 7D probability distributions of the AAMD surface and the ASCG-generated surface is 0.132.

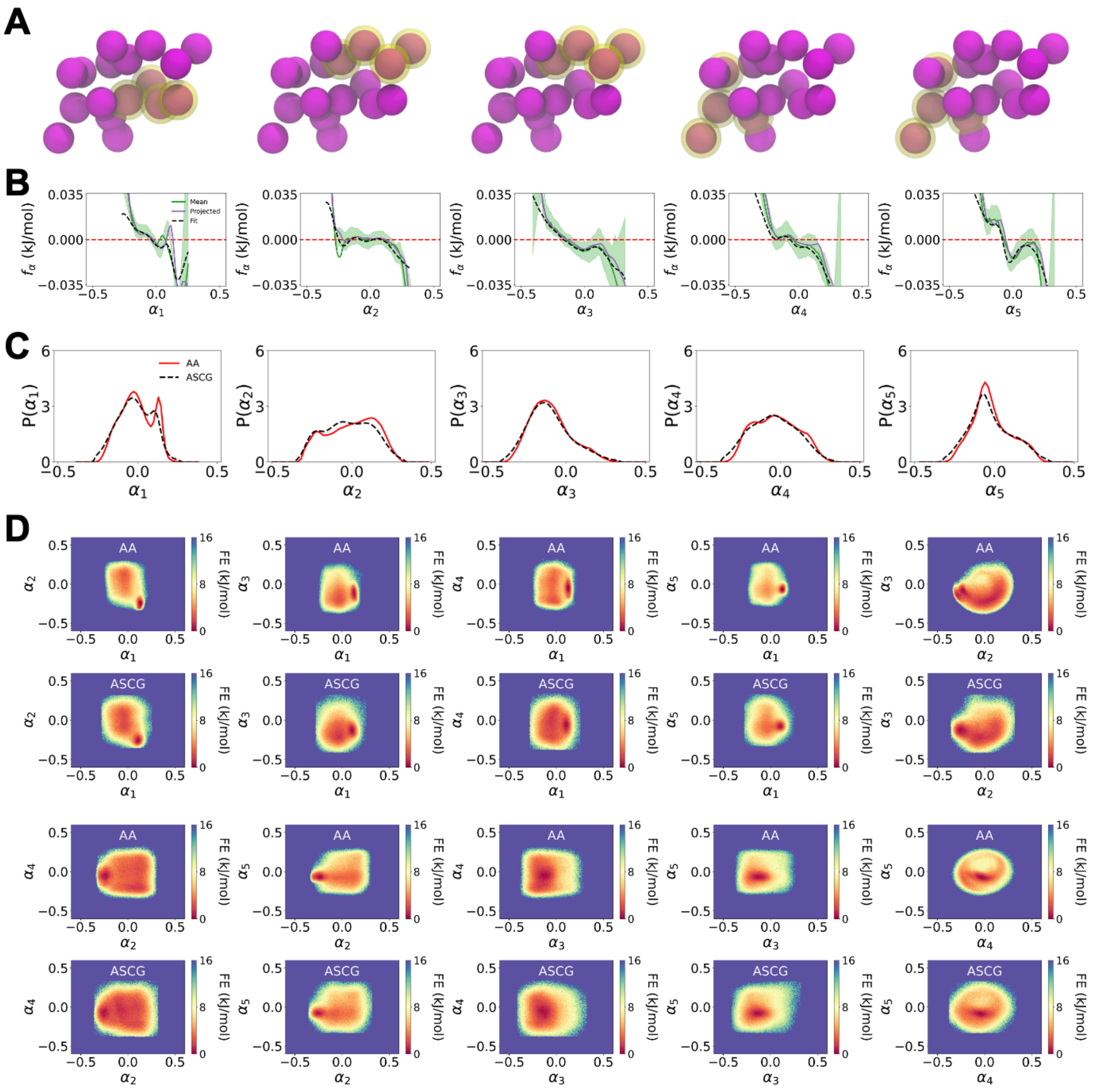


**Fig. S10.** Complete mapping, parametrization, and integration results of 5 ASCG components for chignolin. (A) ASCG mapping of $\alpha_1$ through $\alpha_5$, in order from left to right, depicted in yellow. (B) Force ($f_\alpha$) profiles for $\alpha_1$ through $\alpha_5$. The mean force represents the average forces along each ASCG component after transformation, shown with a 95% confidence interval. Projected forces represent forces estimated by calculating a 1D free energy surface from the 1D probability density of the atomistic data and then taking the negative gradient. Fit forces are predicted forces after fitting a multivariate nearest neighbors algorithm to the averaged ASCG transformed forces. (C) 1D probability distributions along $\alpha_1$ through $\alpha_5$ comparing the atomistic (AA) distribution with the ASCG distribution calculated after integrating $\alpha_1$ through $\alpha_5$. (D) 2D slices of the 5D free energy (FE) surface between each pair of ASCG components, generated from AAMD trajectories (top) and from ASCG integration (bottom). The JSD between the underlying 5D probability distributions of the AAMD surface and the ASCG-generated surface is 0.037.

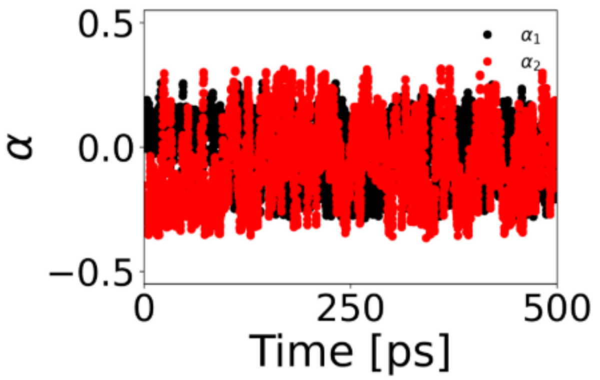


**Fig. S11.** Time series profile of an example ASCG-generated chignolin trajectory, plotting 500 ps of $\alpha_1$ and $\alpha_2$.

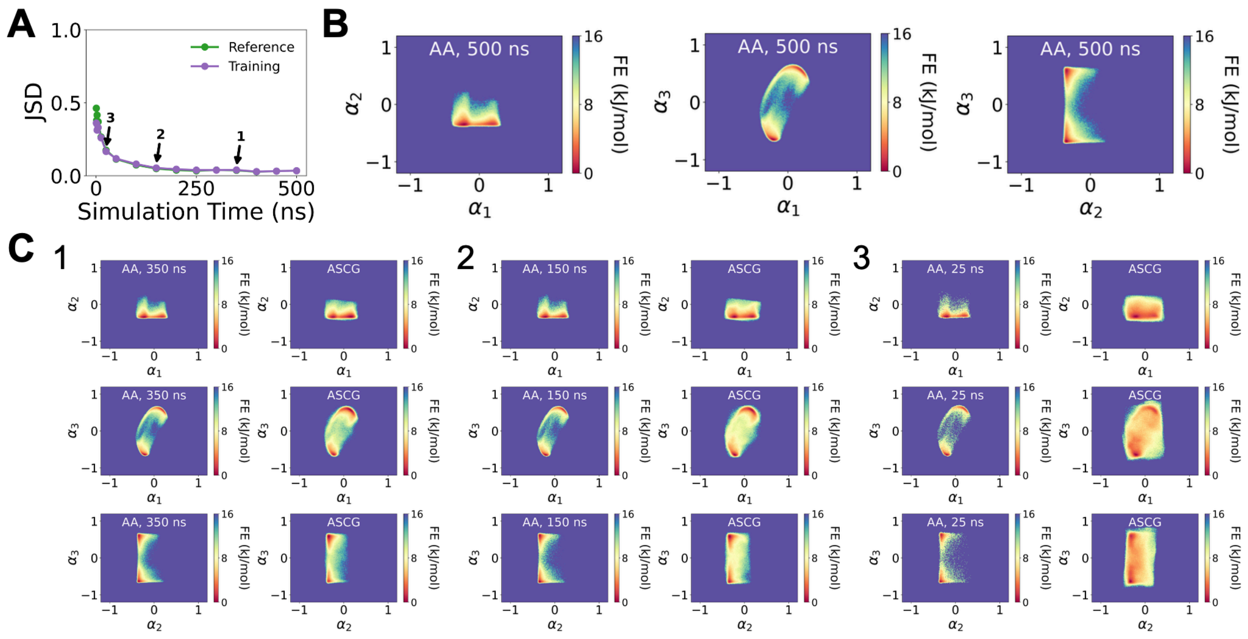


**Fig. S12.** Critical limit of limited data for dialanine. A) The JSD as a function of total simulation time of dialanine used for training. The JSD between the 3D probability distribution of ASCG-generated trajectories is plotted in comparison to the full reference dataset (green) and the shortened dataset that was used to train ASCG (purple). B) The 2D slices of the 3D free energy surface for the full atomistic dataset. C) 2D slices of the 3D free energy surfaces of the shortened atomistic trajectory (left) and of the ASCG-generated trajectory (right) after applying the ASCG method to the shortened trajectory for (1) 350 ns, (2) 150 ns, and (4) 25 ns.

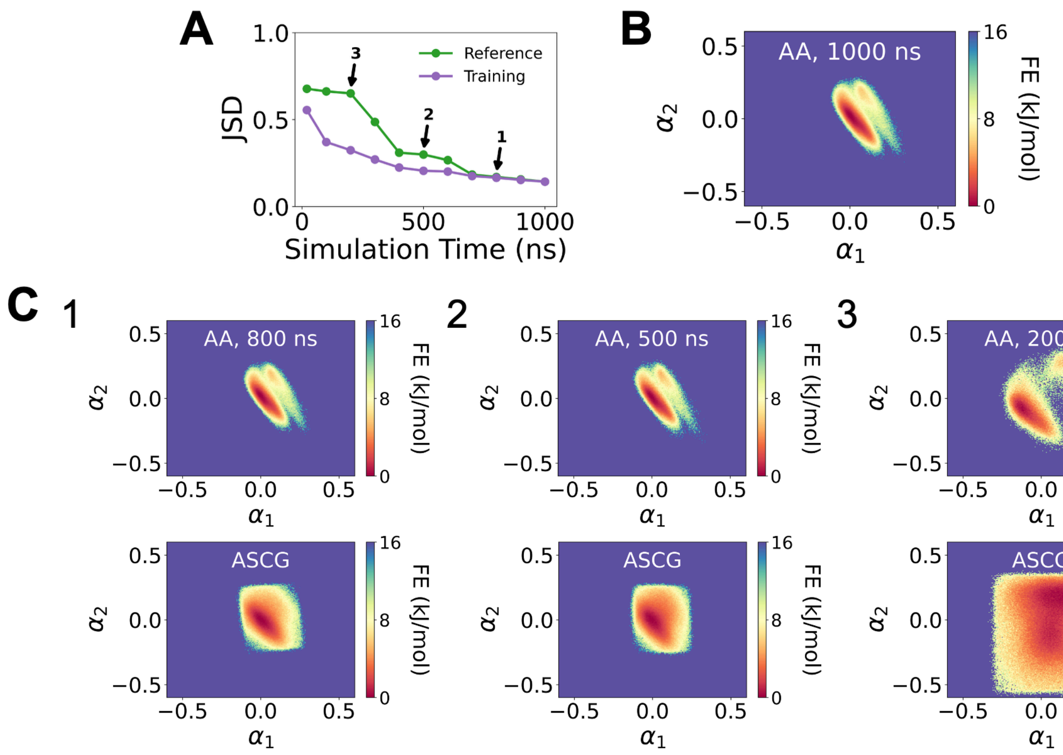


**Fig. S13.** Critical limit of limited data for Trp-cage. A) The JSD as a function of total simulation time of Trp-cage used for training. The JSD between the 7D probability distribution of ASCG-generated trajectories is plotted in comparison to the full reference dataset (green) and the shortened dataset that was used to train ASCG (purple). B) The 2D slice of the free energy surface for the full atomistic dataset of $\alpha_1$ and $\alpha_2$. C) 2D slice of the free energy surfaces of the shortened atomistic trajectory (top) and of the ASCG-generated trajectory (bottom) after applying the ASCG method to the shortened trajectory for (1) 800 ns, (2) 500 ns, and (3) 200 ns.

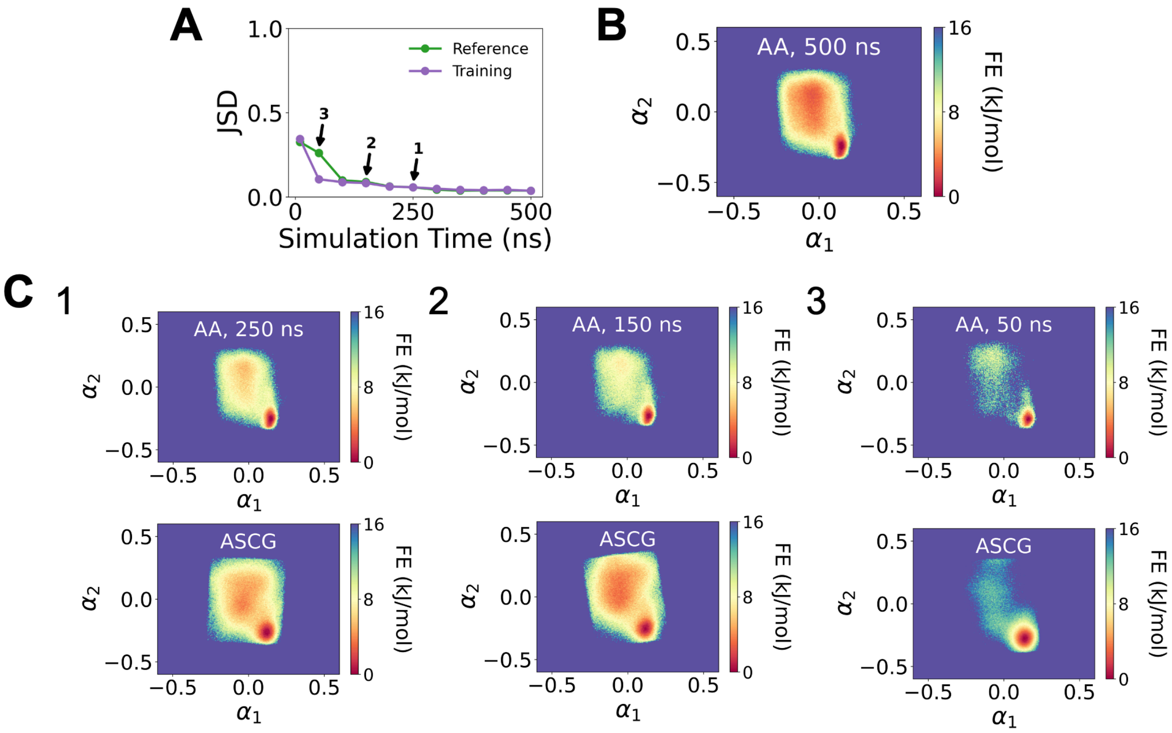


**Fig. S14.** Critical limit of limited data for chignolin. A) The JSD as a function of total simulation time of chignolin used for training. The JSD between the 5D probability distribution of the ASCG-generated trajectories is plotted in comparison to the full reference dataset (green) and the shortened dataset that was used to train ASCG (purple). B) The 2D slice of the 5D free energy surface for the full atomistic dataset of $\alpha_1$ and $\alpha_2$. C) 2D free energy surfaces of the shortened atomistic trajectory (top) and of the ASCG-generated trajectory (bottom) after applying the ASCG method to the shortened trajectory for (1) 250 ns, (2) 150 ns, and (3) 50 ns.

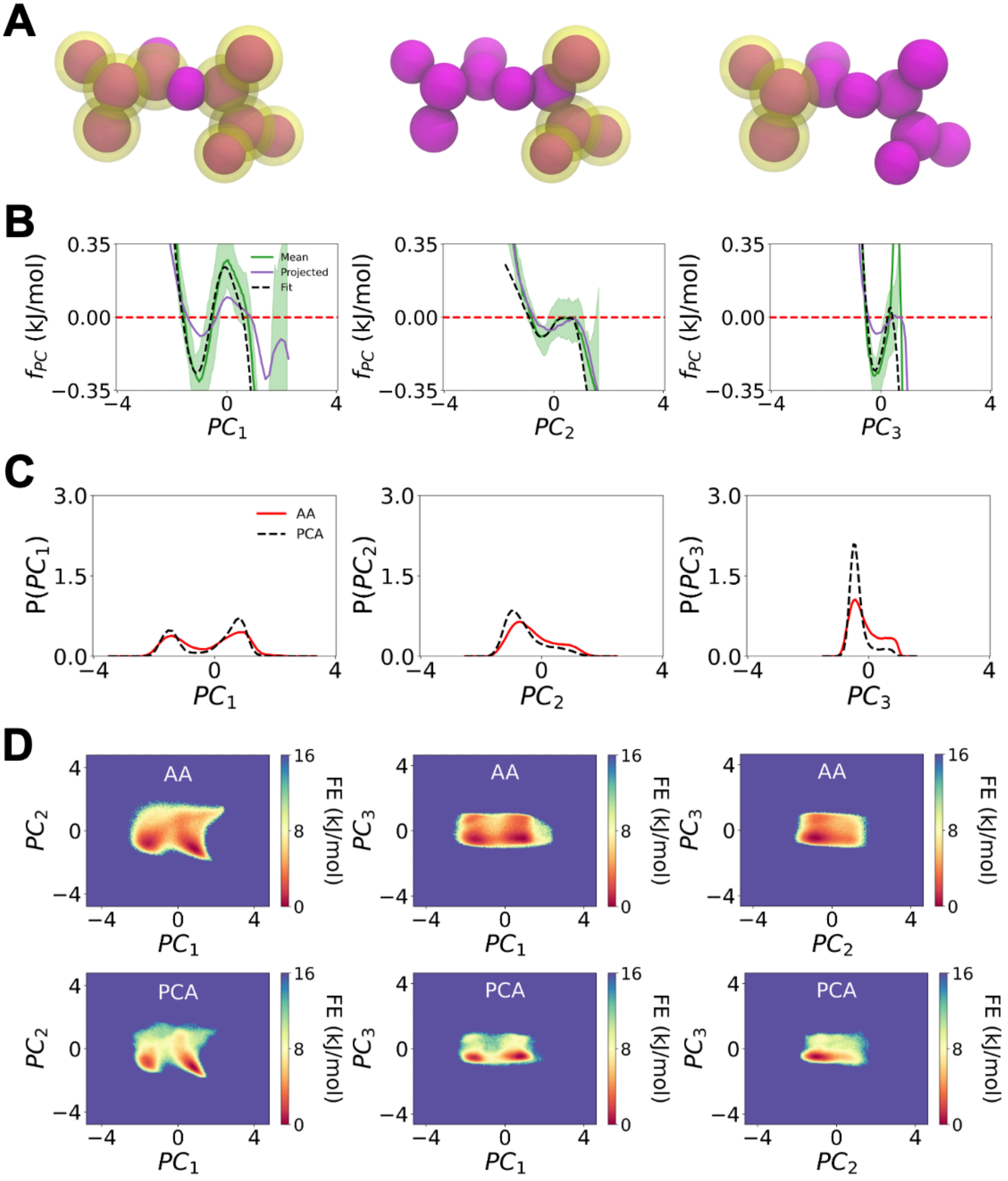


**Fig. S15.** Considering PCA as an alternative unified coarse-graining framework with dialanine with the first 3 PCA components. A) The ASCG mapping of PC1 (left), PC2 (middle), and PC3 (right) is depicted in yellow. B) The mean force represents the average forces along each PC1 (left), PC2 (middle), and PC3 (right) after transformation, shown with a 95% confidence interval. Projected forces represent forces estimated by calculating a 1D free energy surface from the 1D probability density of the atomistic data and then taking the negative gradient. Fit forces are predicted forces after fitting a multivariate nearest neighbors algorithm to the averaged PCA-transformed forces. B) 1D probability distribution for PC1 (left), PC2 (middle), and PC3 (right), and C) 2D slices of the 3D free energy surface of PC1 and PC2 (left), PC1 and PC3 (middle), and PC2 and PC3 (right) generated from atomistic trajectories (top) compared with the 2D slices of the free energy surface generated from ASCG trajectories (bottom). The JSD between the underlying 3D probability distribution of the full-length atomistic trajectory surface and the PCA-generated surface is 0.099.

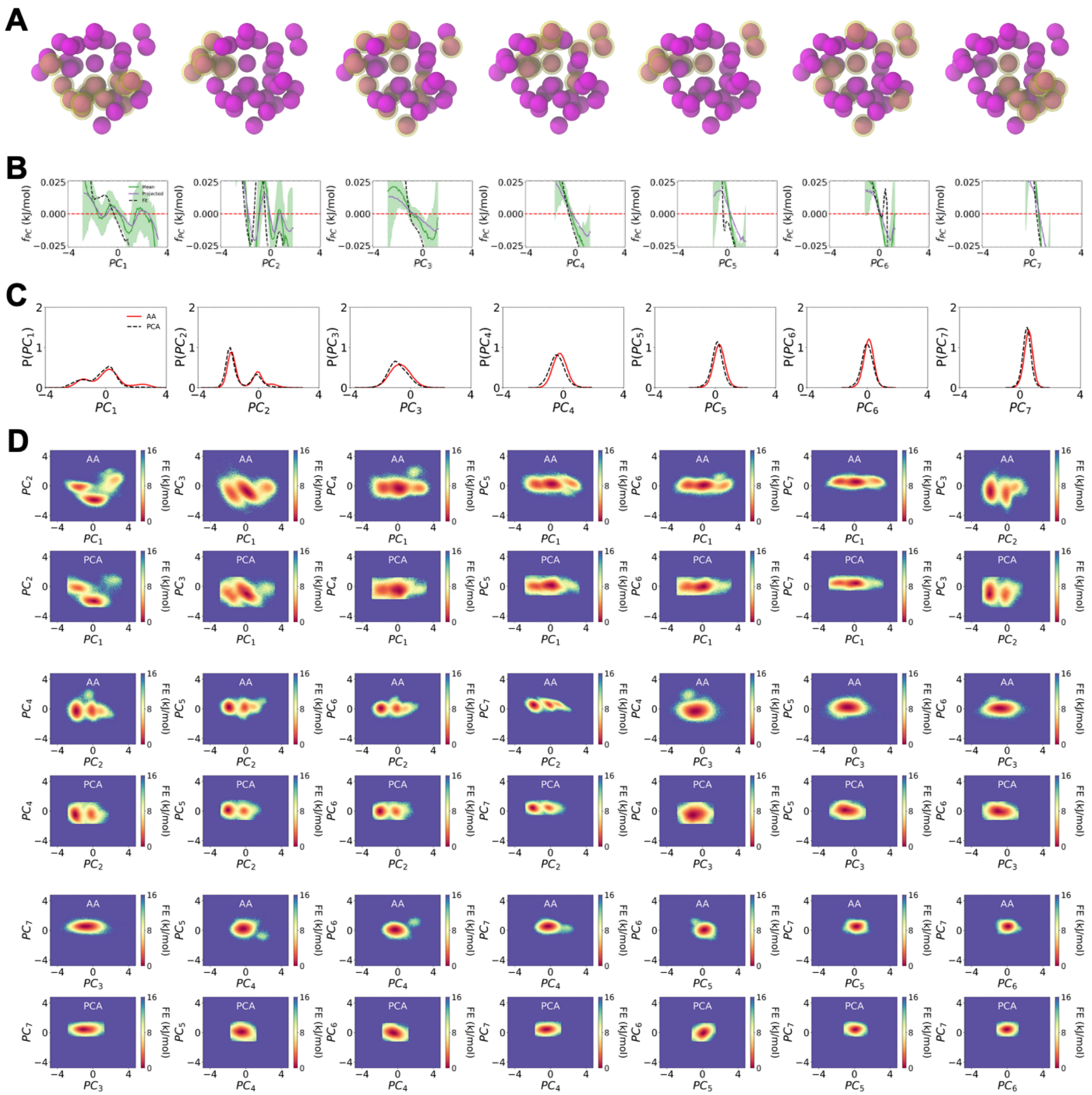


**Fig. S16.** Considering PCA as an alternative unified coarse-graining framework with Trp-cage with the first 7 PCA components. A) The ASCG mapping of PC1 through PC7 in order from left to right is depicted in yellow. B) The mean force represents the average forces along each PC1 through PC7 after transformation, shown with a 95% confidence interval. Projected forces represent forces estimated by calculating a 1D free energy surface from the 1D probability density of the atomistic data and then taking the negative gradient. Fit forces are predicted forces after fitting a multivariate nearest neighbors algorithm to the averaged PCA-transformed forces. B) 1D probability distribution for PC1 through PC7 and C) 2D slices of the 3D free energy surface of each pair of PC components generated from atomistic trajectories (top) compared with the 2D slices of the free energy surface generated from ASCG trajectories (bottom). The JSD between the underlying 7D probability distribution of the full-length atomistic trajectory surface and the PCA-generated surface is 0.244.

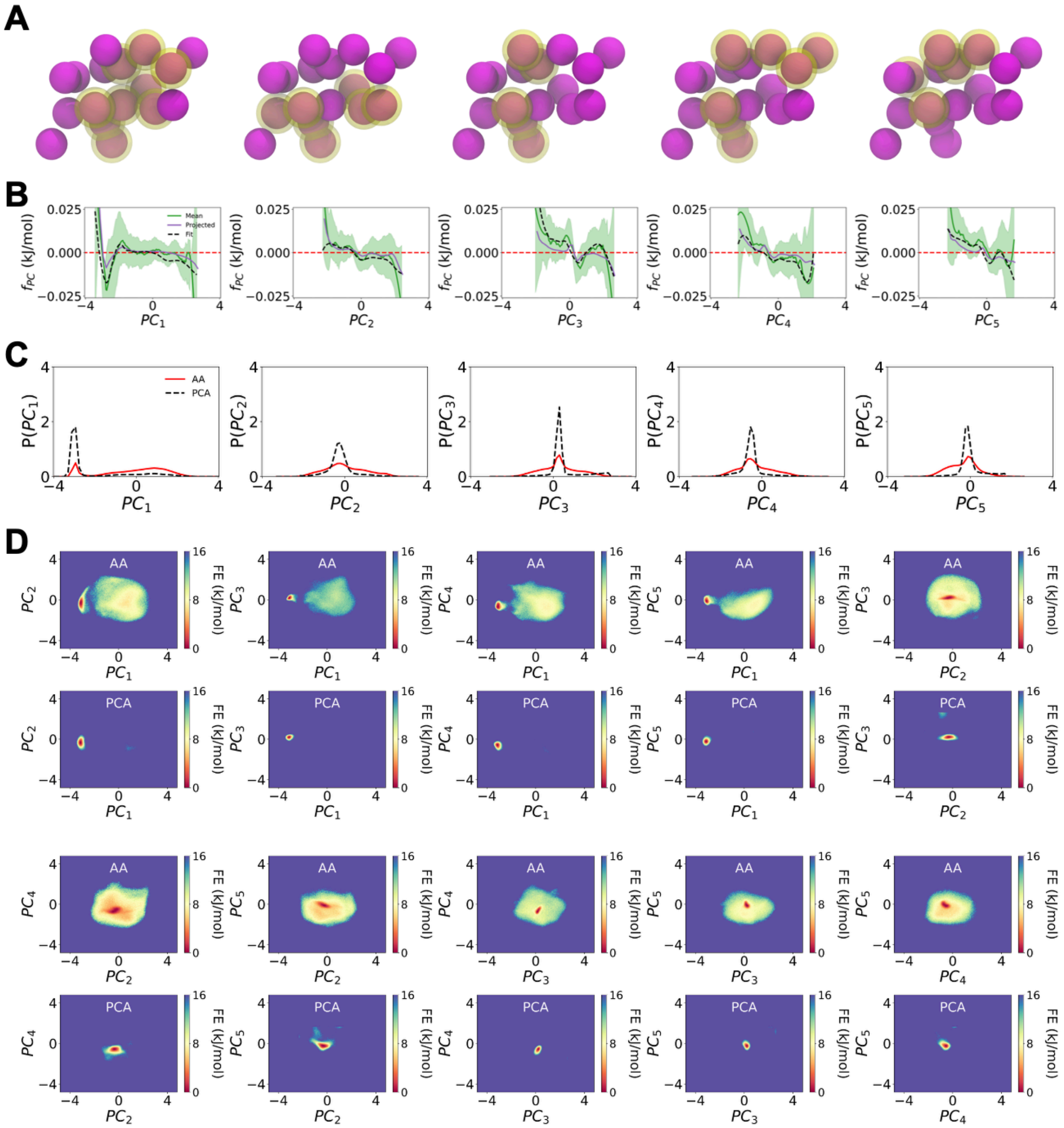

**Fig. S17.** Considering PCA as an alternative unified coarse-graining framework with chignolin with the first 5 PCA components. A) The ASCG mapping of PC1 through PC5, in order from left to right, is depicted in yellow. B) The mean force represents the average forces along each PC1 through PC5 after transformation, shown with a 95% confidence interval. Projected forces represent forces estimated by calculating a 1D free energy surface from the 1D probability density of the atomistic data and then taking the negative gradient. Fit forces are predicted forces after fitting a multivariate nearest neighbors algorithm to the averaged PCA-transformed forces. B) 1D probability distribution for PC1 through PC5 and C) The 2D slices of the free energy surface of each pair combination of PC1 through PC5 generated from atomistic trajectories (top) compared with the 2D slice of the free energy surface generated from ASCG trajectories (bottom). The JSD between the underlying 5D probability distribution of the full-length atomistic trajectory surface and the PCA-generated surface is 0.381.
